# SmartEM: machine-learning guided electron microscopy

**DOI:** 10.1101/2023.10.05.561103

**Authors:** Yaron Meirovitch, Ishaan Singh Chandok, Core Francisco Park, Pavel Potocek, Lu Mi, Shashata Sawmya, Yicong Li, Thomas L. Athey, Vladislav Susoy, Neha Karlupia, Yuelong Wu, Daniel R. Berger, Richard Schalek, Caitlyn A. Bishop, Daniel Xenes, Hannah Martinez, Jordan Matelsky, Brock A. Wester, Hanspeter Pfister, Remco Schoenmakers, Maurice Peemen, Jeff W. Lichtman, Aravinthan D.T. Samuel, Nir Shavit

## Abstract

Connectomics provides nanometer-resolution, synapse-level maps of neural circuits to understand brain activity and behavior. However, few researchers have access to the high-throughput electron microscopes necessary to generate enough data for whole brain or even whole circuit reconstruction. To date, machine-learning methods have been used after the collection of images by electron microscopy (EM) to accelerate and improve neuronal segmentation, synapse reconstruction and other data analysis. With the computational improvements in processing EM images, acquiring EM images has now become the rate-limiting step in automated connectomics. Here, in order to speed up EM imaging, we integrate machine-learning into real-time image acquisition in a single-beam scanning electron microscope. This SmartEM approach allows an electron microscope to perform intelligent, data-aware imaging of specimens. SmartEM saves time by allocating the proper imaging time for each region of interest – first scanning all pixels rapidly, then rescanning more slowly only the small subareas where a higher quality signal is required. We demonstrate that SmartEM achieves up to a ~7-fold acceleration of image acquisition time for connectomic samples using a commercial single-beam SEM in samples from nematodes, mice and human brain. We apply this fast imaging method to reconstruct a portion of mouse cerebral cortex with an accuracy comparable to traditional electron microscopy.

## Introduction

Serial-section Electron Microscopy (ssEM) is widely used to reconstruct circuit wiring diagrams in entire brains of small animals like *C. elegans, Drosophila*, and zebrafish (White et al., 1986; Hildebrand et al., 2017; Witvliet et al., 2021; Winding et al., 2023; Dorkenwald et al., 2024) and brain regions in mammals (Bock et al., 2011; Kasthuri et al., 2015; Morgan et al., 2016; Abbott et al., 2020; Lu et al., 2023; Song et al., 2023). Comparing the growing numbers of connectomes of animals with different genetic backgrounds, life experiences, and diseases will illuminate the anatomical nature of learning, memory, and developmental plasticity, the nature of brain evolution, as well what kinds of anatomical abnormalities cause neuropathology and disease (Kornfeld et al., 2020; Loomba et al., 2022; Karlupia et al., 2023; Bidel et al., 2023; Shapson-Coe et al., 2024). To achieve wide-scale deployment for comparative connectomics, data acquisition and analysis pipelines need to become more widely available (Swanson and Lichtman, 2016). At present, connectome datasets are mostly acquired by the few laboratories and institutions equipped with specialized and expensive high-throughput electron microscopes such as the TEMCA2 (Transmission Electron Microscopy Camera Array 2), GridTape, or the Zeiss 61- or 91-beam scanning electron microscope (SEM) (Zheng et al., 2018; Phelps et al., 2021; Shapson-Coe et al., 2024). Until recently, dataset acquisition had not been a limiting factor in connectomics (Lichtman et al., 2014). A more significant bottleneck had been data analysis – segmenting serial-section electron micrographs to reconstruct the shape and distribution of nerve fibers, identify synapses, and map circuit connectivity. However, recent improvements in machine-learning and image analysis (Beier et al., 2017; Januszewski et al., 2018; Lee et al., 2019; Meirovitch et al., 2019; Sheridan et al., 2023; Popovych et al., 2024) have dramatically sped data analysis, creating a need for faster image acquisition. The field needs more electron microscopes to deliver datasets as fast as they can now be analyzed. One way to meet this need is to enable widely-available electron microscopes, like more affordable single-beam SEMs, to collect connectomic datasets.

When using a single-beam SEM for connectomics, the time budget for image acquisition is currently mostly dictated by the dwell time that the electron beam spends on each pixel. In practice, SEM imaging of well-prepared, high-contrast, electrondense tissue for connectomics usually uses dwell times ⩾ ~1000 ns/pixel. By comparison, the time spent moving the beam between pixels is negligible; many modern SEMs use electrostatic scan generators that can rapidly deflect the electron beam to any pixel in an image (Anderson et al., 2013; Mohammed and Abdullah, 2018). To accelerate an SEM for connectomics, one must reduce the total dwell time over all pixels, but without losing information needed to accurately determine the wiring diagram.

For connectomics, the salient measure of image accuracy is neuronal segmentation – being able to correctly identify each neuron’s border (membrane boundary and extracellular space) and to correctly identify each synapse. In standard SEM, image acquisition is done by specifying a fixed homogeneous dwell time for all pixels. The longer the dwell time, the higher the signal-to-noise per pixel, and the more accurate the segmentation, up to a point at which accuracy saturates. Thus, there is a fundamental trade-off between SEM imaging time and segmentation accuracy; rapid image acquisition can miss critical information that is essential to segmentation. Previous approaches to improving segmentation accuracy with rapidly acquired images have involved post-acquisition image processing such as denoising (Minnen et al., 2021) or “super-resolution”/upsampling (Fang et al., 2021). However, image processing that works entirely after the completion of image acquisition is limited by the amount of original information acquired. No technique unambiguously “creates” information that was not acquired in the first place.

Our solution to the problem of missing information in a rapidly acquired image is to recover information during realtime microscope operation. To do this, we developed a “smart” SEM pipeline that rapidly identifies error-prone regions as well as high-salience regions (such as synapses) in every rapidly acquired image, and then immediately rescans only these regions but now more slowly (longer dwell times per pixel). We define error-prone regions as only those whose rescanning at longer dwell time would confer better segmentation accuracy to a composite image, built from the initial rapidly acquired image (adequate where segmentation is accurate at short dwell time) and targeted rescans (necessary where accuracy is initially insufficient). When error-prone and high-salience regions are relatively few in number and small in size, selective rescanning adds little to the total image acquisition time budget compared to acquiring a uniform long dwell time image, while also restoring segmentation accuracy. We sought an image acquisition pipeline that achieves the accuracy of uniform long dwell time acquisition with nearly the speed of uniform short dwell time acquisition.

We implemented ‘smartness’ in the pipeline with efficient machine-learning algorithms running within SEM computer hardware. This pipeline, called SmartEM, can be applied in any context where images exhibit high spatial heterogeneity in segmentation accuracy as a function of imaging time – a fundamental characteristic of brain images where nerve fibers and synapses can vary in size and density from region to region. Unavoidable spatial heterogeneity in any specimen is why a smart selection of which regions to collect at short dwell times and which regions to rescan at long dwell times can achieve full segmentation accuracy but with less total dwell time. Our experiments with applying SmartEM to three different connectomic datasets using two widely available SEMs yielded between ~5–7× speedup. Because spatial heterogeneity characterizes numerous SEM imaging data, SmartEM can be applied to speed reconstruction of not only other specimens in biology, but also in material sciences and in electronic circuit fabrication.

## Results

### Suitability of adaptive dwell times for connectomics

To establish the rationale for our connectomics pipeline by SEM – automatically applying short dwell times to most areas that are “easy” to segment and long dwell times to fewer areas that are “hard” to segment – we quantitatively tested how spatial heterogeneity in representative mammalian brain images affects segmentation accuracy with different dwell times. To perform these tests, we used a recent high-quality sample comprising 94 sections of mouse visual cortex (Karlupia et al., 2023). We reimaged these 94 sections at 4 nm pixel resolution using a Verios 5 HP SEM from Thermo Fisher Scientific at a range of fixed dwell times from 25 to 1200 ns/pixel.

We note that when these images were originally acquired in a previous study using multi-beam SEM, the dwell time was 800 ns/pixel (Karlupia et al., 2023). This dwell time was determined by an expert operator and is close to the 800–1000 ns/pixel needed for maximal segmentation accuracy for this dataset (**Figure 1A, 1B**).

**Figure 1.**
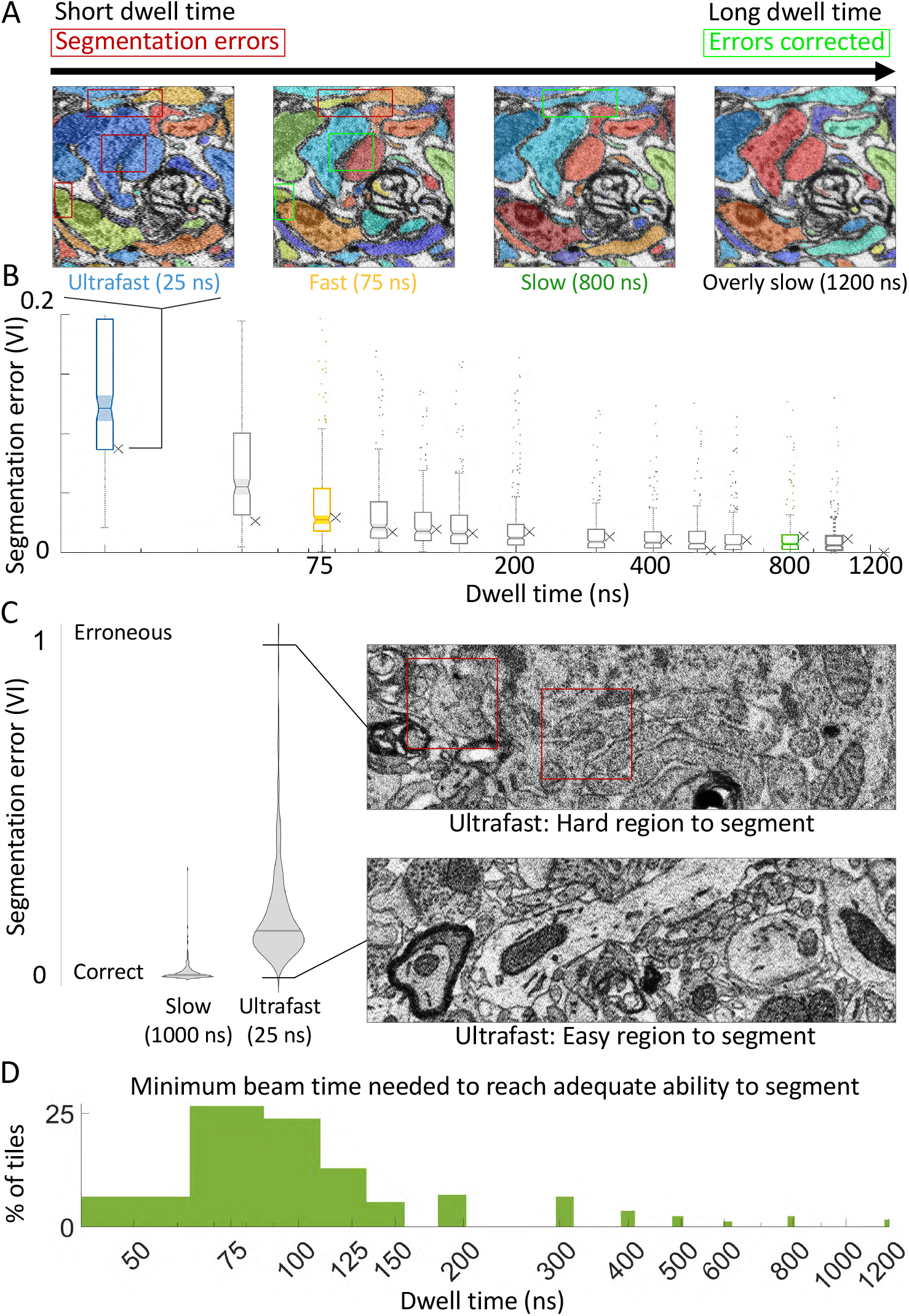
The effect of the beam’s dwell time on the ability to segment the EM into neuronal elements. **A**. Scanning the same EM tile with different dwell times. Short dwell time scans result in segmentation errors (red squares) that are resolved by longer scans (green squares). Increasing the dwell time improves the segmentation accuracy of short dwell time images (25 and 75 ns/pixel) but does not improve the segmentation accuracy of sufficiently long dwell time images (800 ns/pixel). **B**. The segmentation quality of the same images used in (A) are represented by x markers, alongside the distribution of segmentation qualities of 256 images (scatter and boxes) for 13 dwell times, from 25 to 1000 ns/pixel, calculated relative to a reference image taken at 1200 ns/pixel. Segmentation error is quantified by variation of information (y-axis). VI drops rapidly with increased dwell times, saturating with dwell times near 800 ns/pixel. Wide distributions shown by whiskers at each dwell time indicate that some image tiles can be accurately segmented at short dwell times. **C**. Segmentation of neuronal tissue has varying quality due to tissue heterogeneity: taking an image at 25 ns/pixel could lead to an image that can be segmented at high quality (bottom image) or low quality (top image), compared to taking the images slowly (at 1000 ns/pixel). **D**. The majority of image regions (greens areas add up to 1.00) can be segmented at shorter dwell times (75 to 125 ns/pixel), while some regions require longer dwell times (between 400 to 800 ns/pixel) to reach the segmentation quality criterion. Thus, adapting dwell time for different regions would save imaging time without reducing segmentation quality.

Our segmentation algorithm – mapping EM images to border predictions (EM2B) followed by a standard watershed transform – provided an objective assessment of segmentation accuracy of images collected with different dwell times. We adapted EM2B to SEM images taken with different dwell times (see **Supplementary Information**). We automatically segmented 256 randomly selected 2048 × 1768 pixel regions taken from the 94-section sample imaged at 14 different dwell times. Automatic segmentation with ultrafast dwell times (25 ns/pixel) produced frequent merge and split errors compared to automatic segmentation of the same regions with overly slow dwell times (1200 ns/pixel) (**Figure 1A**). As dwell times increased, segmentation errors gradually disappeared.

To quantify segmentation accuracy, we calculated the Variation of Information (VI; Meila (2003)) between each automatically segmented region at each shorter dwell time and the segmentation obtained at the longest dwell time (**Figure 1B**). Segmentation accuracy increased with longer dwell times, and saturated at 800-1000 ns/pixel, consistent with the rule-of-thumb practice in choosing the dwell times for connectomics. At 25 ns/pixel, acquisition speed is 40× faster than at 1000 ns/pixel, but with lower segmentation accuracy.

Brain tissue is typically heterogeneous, with some image regions easier and others harder to segment accurately (**Figure 1C, 1D**). Thus, segmentation accuracy varied substantially from region to region. For slow dwell times (1000 ns/pixel), segmentation accuracy was narrowly distributed around small VI, indicating fewer segmentation errors. For ultrafast dwell times (25 ns/pixel), segmentation accuracy was broadly distributed. Some regions exhibited the same low VI with both ultrafast and slow dwell times (“easy” to segment regions). In contrast, some regions exhibited drastically higher VI for ultrafast dwell times than slow dwell times (“hard” to segment regions) (**Figure 1C**). For each region, we determined the minimum dwell time to reach the same segmentation accuracy as that produced by the longest dwell time (see **Supplementary Information: Determination of maximal segmentation quality**). We observed a broad distribution of minimum dwell times across pixel regions in this mouse cortex sample. Most 2048 × 1768 pixel regions are accurately segmented with dwell times *≤*150 ns/pixel, but some (~25%) required longer dwell times. Minimum dwell times exhibited a broad-tailed distribution from 50-1200 ns/pixel (**Figure 1D**).

### Challenges in implementing SmartEM

We sought a SmartEM pipeline that could both identify and adapt to the spatial heterogeneity in the segmentation accuracy of brain tissue at various dwell times. Implementing such a pipeline on an SEM poses several challenges. First, the SEM needs to automatically identify regions likely to produce segmentation errors when acquired rapidly. Next, the SEM needs to slowly rescan pixel neighborhoods within and around these “error-prone” regions to improve segmentation. Finally, the pipeline needs to accurately segment composite images built from the initial rapidly acquired images fused with rescanned error-prone regions. Below, we describe the solutions to these challenges that form our SmartEM pipeline.

### Detecting error-prone regions by an SEM

To identify error-prone regions in rapidly acquired images, we developed a machine learning (ML) algorithm to run on the microscope’s support computer. To take a particular example (**Figure 2A**), a short dwell time scanned image produced a segmentation merge error (see red arrow) that would have been avoided with a longer dwell time scan of the same tile (middle panel). By use of a neural network (ERRNET; see below), it was possible to identify the error-causing location in the rapidly acquired image and specify the error-prone region to be rescanned that would remedy segmentation errors (yellow outline in middle and right panels). This region includes the poorly defined cell membranes causing the merge error (red line in right panel). ERRNET can operate in real-time within the SEM computer when equipped with a commodity GPU. This network runs faster than the short scan image acquisition (<100 ns/pixel) and can be further sped up by parallelization with multiple GPUs. A related idea where EM acquisition is guided based on uncertainty measures estimated by neural network models was described in Shavit et al. (2021).

**Figure 2.**
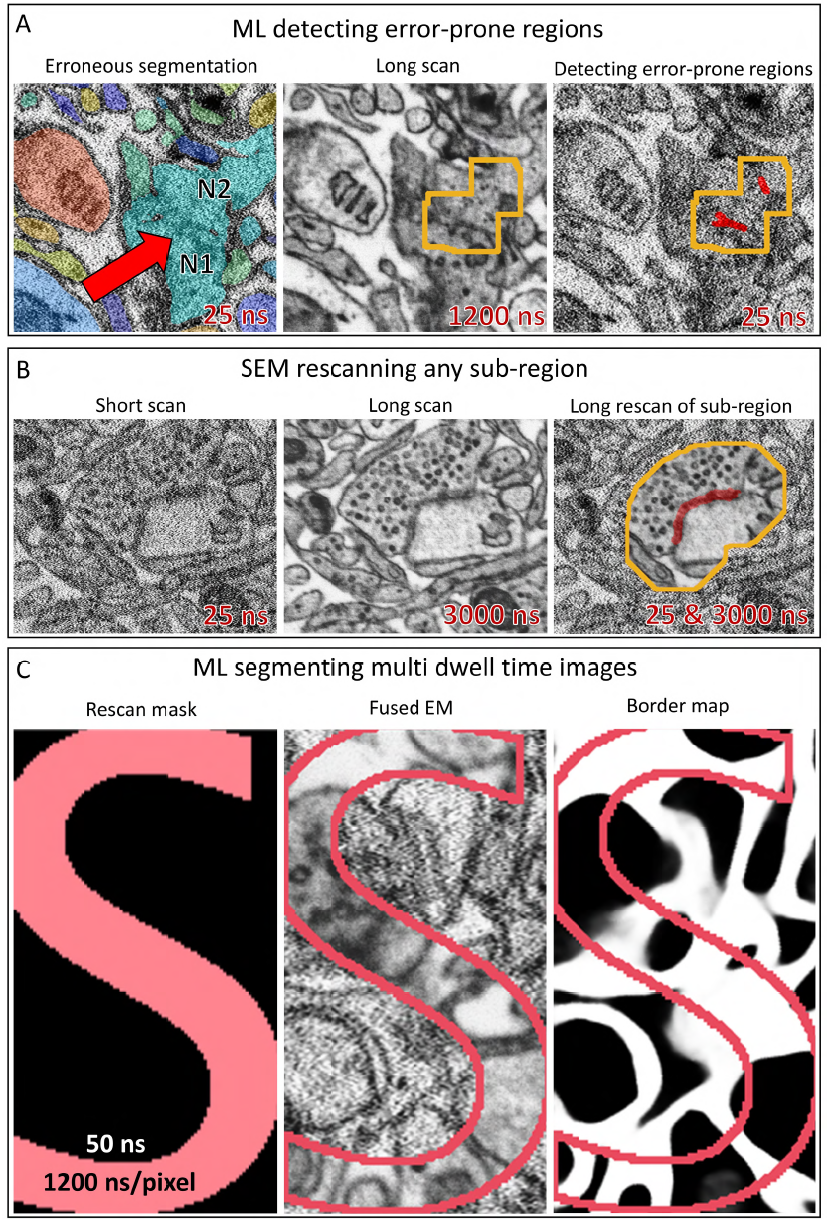
Smart microscope challenges. **A**. An erroneous segmentation of a rapidly acquired image (25 ns/pixel) with a red arrow indicating the location of a merge error between two neurons (N1, N2). Acquiring the same image at a long dwell time of 1200 ns/pixel enhances the neuronal boundary (middle). The output of the ERRNET neural network that was trained to predict segmentation errors from EM is shown on the right in red. The yellow outline is a window around the predicted error to provide further context needed for downstream correction. **B**. The SEM readily captures any part of an image at different dwell times, homogeneously at short dwell times (left), homogeneously at long dwell times (middle), or homogeneously at short dwell times with a sub-region taken at long dwell times (right). Here, the yellow outline for the long dwell time sub-region contains a synaptic cleft. **C**. Predicting neuronal borders from fused EM images using FUSEDEM2B.

### Rescanning targeted sub-regions

To use the prediction of error-prone regions during real-time SEM operation, we modified the scanning procedure of the microscope to rescan error-prone regions at long dwell times right after the short scan. In addition to rescanning error-prone regions, neural networks can be trained for data-aware rescan of additional regions of interest like synaptic clefts for applications in connectomics. **Figure 2B** depicts data-aware rescan where the microscope is guided to re-take regions around synaptic clefts that are predicted from an initial short scan image of a section of mammalian cortex. SEMs with electrostatic scan generators are able to conduct efficient and rapid rescan without wasted time in moving the electron beam (Mohammed and Abdullah, 2018; Anderson et al., 2013). When ERRNET and rescan software are seamlessly integrated within SEM computer hardware, the total time spent acquiring an image is the total number of pixels × the short initial dwell time plus the total number of *rescanned* pixels × their long dwell time.

### Segmentation of composite dwell time images

After image acquisition, a smart microscopy pipeline generates a complete rapidly acquired image and set of rescanned regions of each sample acquired at longer dwell times. Composite images are produced by substituting pixels from rescanned regions into corresponding locations in the initially rapidly acquired images, resulting in images with pixels of multiple dwell times. Previous segmentation algorithms for connectomics have dealt with a single pre-fixed dwell time (Lee et al., 2017; Januszewski et al., 2018; Meirovitch et al., 2019; Sheridan et al., 2023) – these algorithms generalize poorly to homogeneous images taken at different dwell times or to heterogeneous images composed of regions taken at different dwell times. The smart microscopy pipeline demands new algorithms to accurately segment composite images where different regions are obtained at different dwell times. We developed a data augmentation training procedure technique for a neural network with a U-Net (Ronneberger et al., 2015) architecture (FUSEDEM2B) to accurately detect borders in an image with heterogeneous dwell times as well as if the image was taken with a single uniformly applied dwell time (see **Supplementary Information**). **Figure 2C** shows an example of an image that has multiple dwell times (long dwell time scanning arbitrarily within an S-shaped region surrounded by short scanning). The predicted borders by FUSEDEM2B are unperturbed when crossing between regions taken with different dwell times.

Thus, the challenges in building a smart microscopy pipeline are met by extensively using machine learning in both guiding image acquisition and image analysis.

### The smart microscopy pipeline

We developed an integrated smart pipeline that meets the above challenges. **Figure 3** illustrates how the pipeline operates on a small tile from the mouse cortex dataset (Karlupia et al., 2023). The core components of SmartEM are outlined in **Figure 4** and their design and implementation are described below in detail.

**Figure 3.**
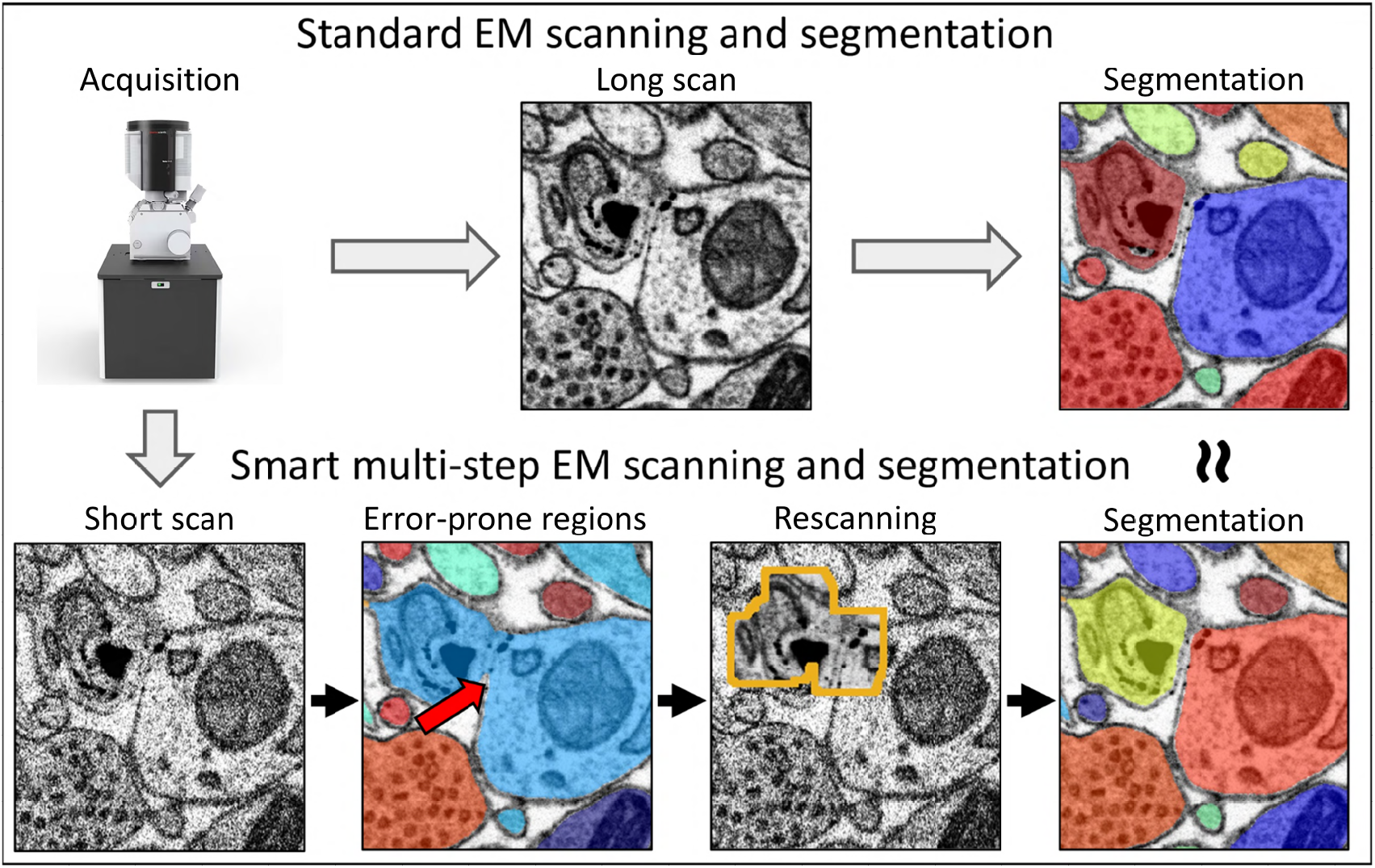
The smart multi-step imaging compared to standard imaging. In standard EM, the sample is first scanned with a long dwell time and then segmented (top). In the SmartEM pipeline, the sample is first scanned at a short dwell time, error-prone regions are detected and rescanned and then segmented.

**Figure 4.**
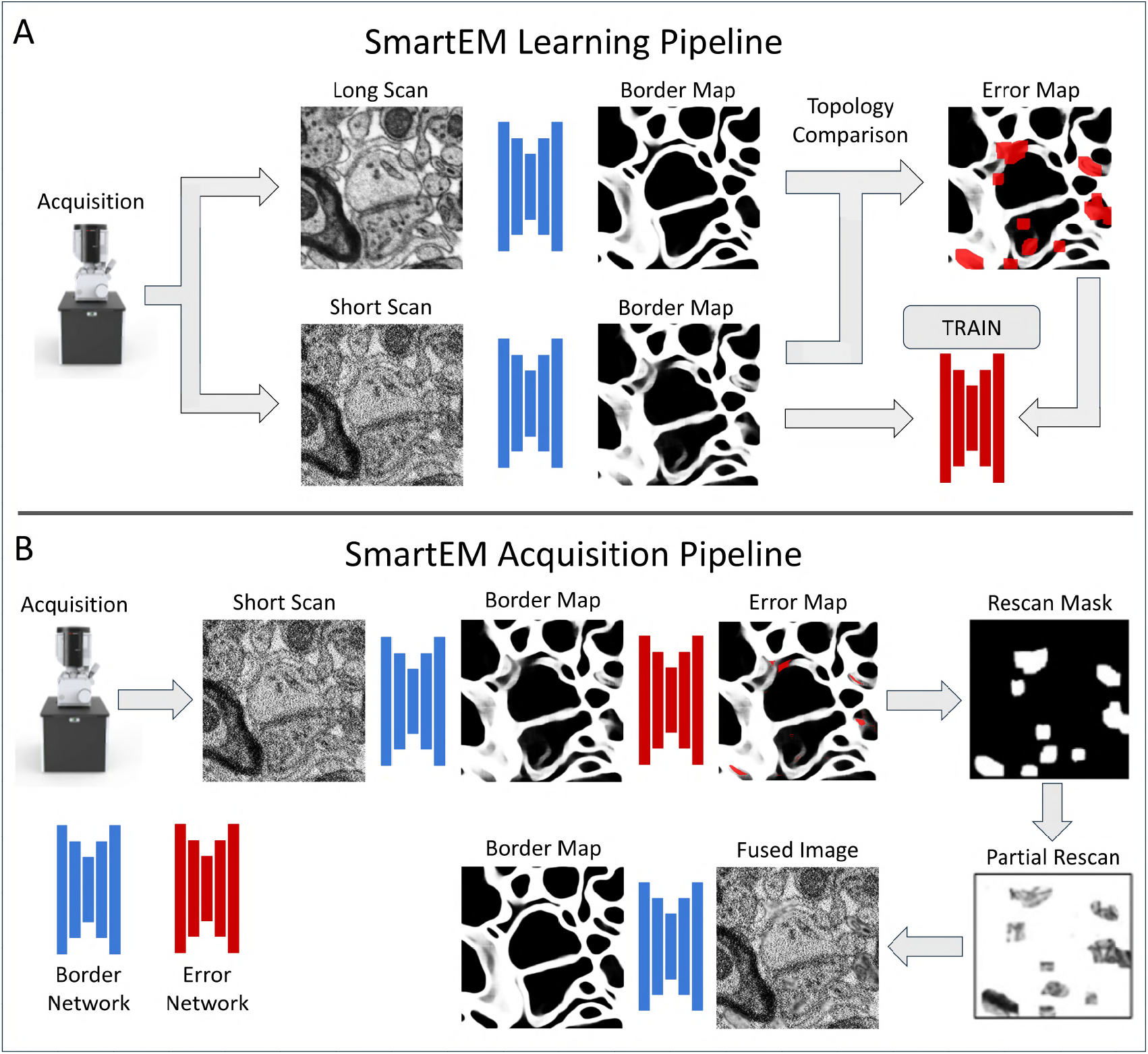
The learning and acquisition phases of SmartEM. **A**. For training, SmartEM requires aligned stacks of high-quality (long scan) images and low-quality (short scan) images. A border detector, FUSEDEM2B (blue), is trained on this dataset to re-produce the high quality results of a border detector that runs only on the long scan images. Once FUSEDEM2B is trained, the border predictions between the short and long dwell times is compared (topology comparison) and a binary error map featuring the differences between the two predictions is produced. A second network, ERRNET (red) is trained to predict this error map from the border predictions of the short dwell time images. **B**. For acquisition, SmartEM first performs a short scan. The trained networks FUSEDEM2B and ERRNET are used to obtain a rescan mask. This region is rescanned at a longer dwell time, resulting in a fused EM image with better segmentation quality.

### Determining the standard dwell time needed for high accuracy segmentation

The goal of the SmartEM pipeline is to reach the same segmentation accuracy as a standard SEM when using a uniform long dwell time scanning regime, but acquiring the images in much less time. To accurately assess the improvement of SmartEM over a standard SEM imaging regime, we needed first to determine the shortest uniform dwell time that leads to accurate segmentation (e.g., 800-1000 ns/pixel in the example in **Figure 1B**).

To accomplish this, we trained a neural network called SLOWEM2B to perform automatic border prediction in long dwell time-acquired images. We collected a diverse subset of long dwell time images from random locations in a specimen, typically twenty 5 × 5 *µ*m^2^ tiles, and performed manual segmentation by an expert to create training data for SLOWEM2B. Next, we used SLOWEM2B to train a different neural network called EM2B that was capable of predicting borders with either long or short dwell time images. Because the SEM allowed for re-imaging the same regions at different dwell times, it was possible to guide the microscope to collect a large sample of EM images from different random locations in the specimen, using different dwell times ranging from 25 to 2500 ns/pixel as shown in **Figure 3**. SLOWEM2B was applied to the long dwell time image at each location to automatically create segmentations that we could use as “ground truth” to train EM2B to predict segmentations in both long and short dwell time images. Both SLOWEM2B and EM2B were implemented using a U-Net architecture.

SLOWEM2B and EM2B calculated the trade-off between pixel dwell time and segmentation accuracy. EM2B was used to automatically segment all dwell time images (e.g. from 25 to 1000 ns/pixel for the mouse cortex dataset) and compare them to a reference automatic segmentation corresponding to the longest dwell time image (e.g. 1200 ns/pixel image). Thus, it was possible to identify the shortest dwell time for which mean accuracy across tiles was not further improved by longer dwell time imaging. This minimum dwell time was defined by SmartEM as the required dwell time to achieve agreement with the longest dwell time segmentation. For the mouse cortex dataset, the minimal dwell time required to achieve maximal segmentation accuracy at 4 nm per pixel was 800–1000 ns/pixel.

### Learning to detect error-causing locations in short dwell time images

To further reduce imaging time we adjusted the pixel dwell time locally based on maintaining segmentation accuracy. Most image regions can be segmented with full accuracy after scanning with a short dwell time. Additional dwell time was only selected for those regions that required longer imaging to segment properly. This selection was accomplished via a neural network (ERRNET) that learned what regions required a longer dwell time after scanning whole images with a short dwell time. ERRNET learns the features of error-causing locations in raw short dwell time images that produce segmentation differences – erroneous merges or splits – in comparison to long dwell time-acquired images.

To assemble “ground-truth” to train ERRNET, the microscope first takes a large set of images from random locations in the specimen at multiple dwell times (e.g. from 25 to 1200 ns/pixel). These images are segmented to distinctly label every contiguous neuron cross section. Automatic labeling can be done using border probabilities, a seeding procedure, and a standard region-growing algorithm such as watershed (Vincent and Soille, 1991). Segmented images at all dwell times are compared to reference segmented images taken with the longest dwell time (1200 ns/pixel for the mouse cortex dataset in **Figures 1A, 1B**, longer than needed for fully accurate segmentation with SLOWEM2B). To automatically learn segmentation discrepancies between short and long dwell time images, we developed a method to produce a binary error mask that defines the morphological differences between two segmented images based on the variation of information (VI) clustering metric (Meila, 2003) (see **Supplementary Information**). We trained ERRNET to predict error-causing regions in short dwell time image as shown in **Figure 4A**. We used the VI metric to detect objects that are morphologically different between segmentations of short and long dwell time images, and then mapped the borders that differ for these objects (described in **Supplementary Information**) (Meila, 2003). We noted that all segmentation errors in short dwell time images can be repaired (i.e. leading to identical segmentation as long dwell time images) by selectively replacing only regions surrounding discrepancy-causing locations in short dwell time images with corresponding regions taken from long dwell time images.

In real-time operation, the SEM must take an initial rapidly acquired image, execute ERRNET to detect error-prone locations, define a rescan mask by padding error-prone locations to capture enough context to improve segmentation accuracy, and then immediately rescan all error-prone regions using longer dwell times (**Figure 4B**).

### Enhancing composite images for improved human readability

The final output of the pipeline are images where sets of small regions are captured with longer dwell times than the rest of the image. Although the raw appearance of short dwell time regions (high pixel noise) and long dwell time regions (low pixel noise) does not degrade segmentation accuracy, it does create unappealing contrasts for human vision. To improve the SmartEM image for human interpretation, we also built an algorithm that translates the style of the SmartEM images to look like standard EM images with homogeneous dwell times. A similar technique was described in Shavit et al. (2021, 2023). This stylized output does not supplant, but is saved in addition to, the raw composite SmartEM images. We note that stylized images often retain the correct details of the ultrastructure seen in homogeneous long dwell time images (**Figure S9**).

### Technique Evaluation

We developed our SmartEM pipeline to expedite connectomics reconstruction on two widely available SEMs, the Verios 5 HP and the Magellan 400L, both from Thermo Fisher Scientific. We quantitatively estimate the practical improvement in quality and speed of this pipeline for connectomics in a variety of tissues, including re-imaging a previously studied mouse cortex (Karlupia et al., 2023), a previously studied human temporal lobe H01 dataset (Shapson-Coe et al., 2024), and a newly prepared male *C. elegans* dataset.

### Improving accuracy

One premise of the smart microscopy pipeline is that automatically detecting error-prone regions and replacing them with longer dwell time pixels will reduce segmentation errors. To test this premise, we compared the accuracy of a segmentation pipeline trained to deal with short dwell time images (FASTEM2B at 100 ns/pixel) to a SmartEM pipeline trained to deal with composite images made from short and long dwell times (FUSEDEM2B at 100 ns/pixel and 2500 ns/pixel). The performance of these networks was compared to the standard segmentation pipeline with long dwell time image acquisition (SLOWEM2B at 2500 ns/pixel). For fair comparison, we used the same long dwell time for the rescanning in the SmartEM pipeline and for the uniform scan in the standard pipeline. We found that using these dwell times, SmartEM is approximately 5× faster than the standard segmentation pipeline with long dwell time image acquisition and approximately 2-3× more accurate (based on VI) than the standard pipeline operating quickly (100 ns/pixel) (**Figure S1**). Thus, fusing long dwell time pixels into a rapidly acquired image can improve segmentation accuracy.

Another premise of the SmartEM pipeline is that the additional time devoted to rescanning part of an image yields a greater improvement in segmentation accuracy than distributing the same extra time across all pixels with a uniform dwell time as shown in **Figure 5A**. To test this premise, we used a “standard” pipeline with competitively fast settings – 400 ns/pixel for *C. elegans* and 75 ns/pixel for the mouse and human cortex datasets. We then compared these images to a SmartEM pipeline configured to match the same overall acquisition time by combining an initial short scan and a targeted longer rescan (**Figure S15**). For the three datasets, the initial SmartEM dwell time was set to 300, 50, and 50 ns/pixel, and the rescan dwell time was set to 800, 150, and 300 ns/pixel, respectively. In each case, we adaptively selected 12.5%, 16.7%, and 8.33% of the most error-prone regions for rescanning to ensure that total acquisition time matched that of the standard pipeline. The procedure for selecting these SmartEM parameters for imaging is described below. We compared the Variation of Information (VI) from 123, 219, and 62 segmented image tiles of each pipeline to reference images taken at a long dwell time, and found that SmartEM produced significantly fewer errors than the standard pipeline (sign tests and distributions of VI differences are in **Figure S15**).

**Figure 5.**
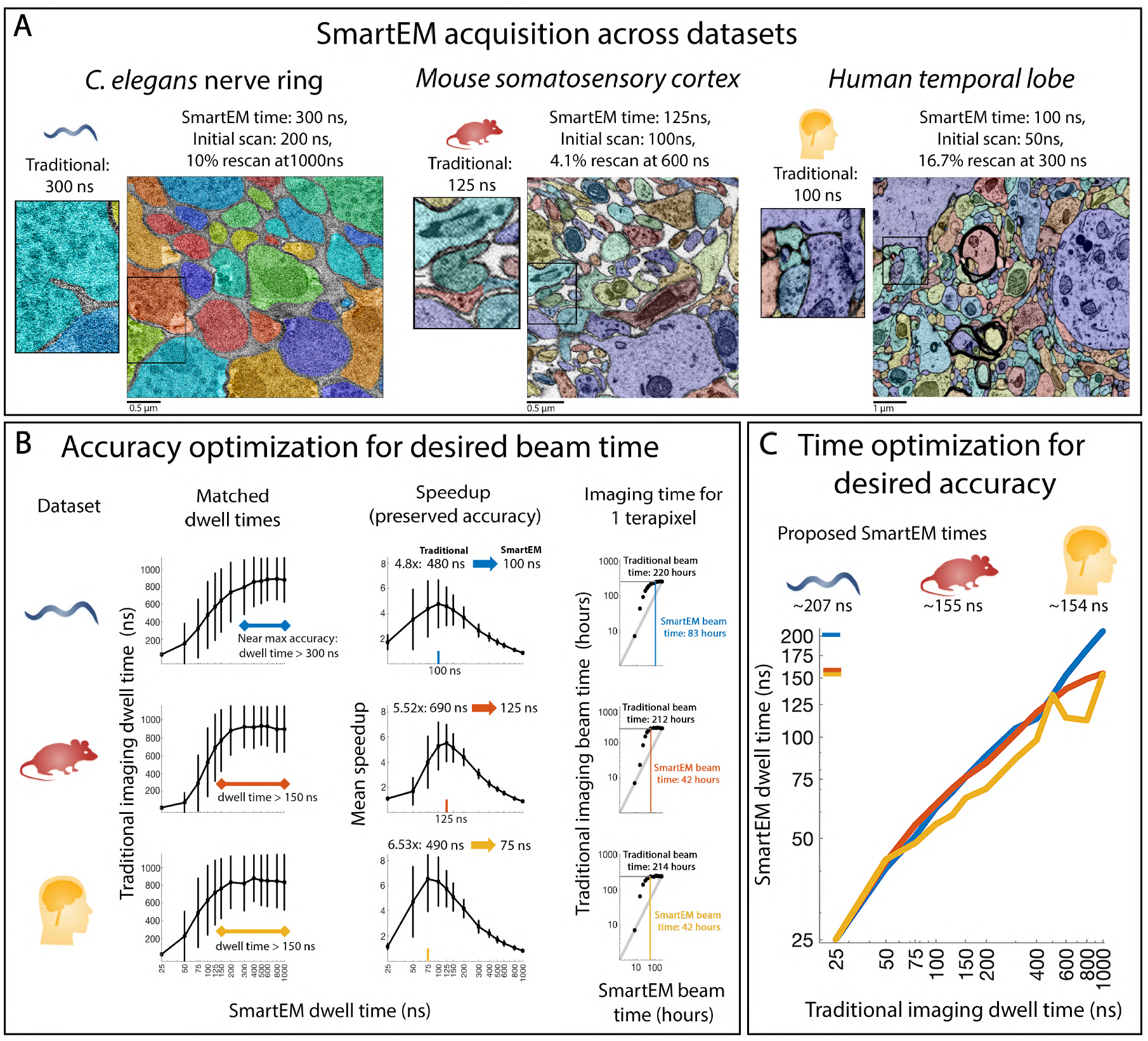
SmartEM acquisition time. **A**. Examples of SmartEM acquisition across three datasets, *C. elegans* nerve ring, mouse somatosensory cortex, and human temporal lobe, visually compared to traditional EM imaging at the same average dwell time. In the first imaging scenario (**B**), the imaging time is constrained by a fixed time budget, which, given a volume size and pixel resolution, determines the average dwell time (beam time per pixel). The task is to identify SmartEM parameters (initial dwell time, rescan dwell time and rescan rate) that optimize segmentation accuracy. **B-left**. For each targeted effective (SmartEM) dwell time (x-axis), we compute the optimal SmartEM parameters and determine the corresponding standard (homogeneous) dwell time (y-axis) required to achieve the same segmentation accuracy. As the SmartEM dwell time increases, the equivalent homogeneous dwell time also rises, approaching an asymptote near 300 ns/pixel (*C. elegans*), 150 ns/pixel (mouse), and 150 ns/pixel (human), corresponding to homogeneous dwell times of 800–1000 ns/pixel in all three datasets. Error bars represent *±*1 s.d. **B-middle**. The resulting speedup (ratio of the homogeneous dwell time to the SmartEM dwell time) from B-left. Maximum speedups occur near the inflection point in B-left, around 100 ns/pixel (*C. elegans*), 125 ns/pixel (mouse), and 75 ns/pixel (human). Increasing the effective SmartEM dwell time beyond these values continues to improve segmentation accuracy but yields diminishing returns in speedup. **B-right**. The data from B-left and B-middle illustrated for a fixed volume of 1 TB at 4 nm/pixel with a slice thickness of 30 nm. **C**. In the second imaging scenario, the desired EM quality is set by a standard pipeline’s dwell time, and the goal is to identify SmartEM parameters that achieve equivalent segmentation quality in minimal imaging time. Near-maximal segmentation quality (comparable to homogeneous 1000 ns/pixel scanning) is attained at roughly 207 ns/pixel (*C. elegans*, blue tick), 155 ns/pixel (mouse, red tick), and 154 ns/pixel (human, orange tick).

### Estimating speedup

We considered two scenarios for the largescale collection of a connectome dataset. The first involves a fixed imaging time budget to acquire a selected data volume at the selected pixel resolution. Here, the task is to intelligently allocate a fixed imaging time to optimize segmentation accuracy.

The second scenario involves matching a fixed image quality to acquire a volume. Here, the task is to determine SmartEM parameters (initial dwell time, rescan dwell time and rescan rate) that maintain the quality of a given traditional dwell time while minimizing the required imaging time. Below we analyze both scenarios.

#### Scenario 1: Optimized accuracy with a fixed imaging time budget

We fix the total imaging time budget for a given specimen. From this requirement, the pixel dwell time is determined after subtracting overhead factors (such as image focusing, astigmatism correction, and mechanical stage movement) from the total budget. For example, the user might need to image a given specimen – 100 × 100 × 100 *µ*m^3^ tissue, cut in 30 nm thick sections, imaged at 4 nm spatial resolution – within 5 days of continuous EM operation. These constraints determine the average dwell time per pixel

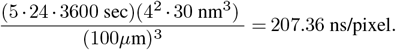

For a standard EM pipeline, 207.36 ns/pixel becomes the homogeneous pixel dwell time. For the SmartEM pipeline, the initial scan and rescan of all error-prone regions should sum to an average of 207.36 ns/pixel. This average dwell time, which we call *effective dwell time*, can be achieved with different combinations of initial dwell time, rescan dwell time, and percentage of rescanned pixels:

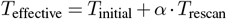

where *T* represents dwell times.

For example, an effective average dwell time of 207.6 ns/pixel is achieved with an initial dwell time of *T*_initial_=100 ns/pixel, rescan rate of *α*=5%, and rescan dwell time of *T*_initial_=(207.36 − 100)*/*0.05=2147.2 ns/pixel. These parameter settings correspond to a specific segmentation accuracy (VI) relative to the reference homogeneous long scan image. SmartEM considers a grid of parameter settings and calculates the *T*_initial_, *T*_rescan_ and *α* settings that produce maximal accuracy (minimal VI) compared to the segmentation of reference tiles, while guaranteeing the effective dwell time (see **Supplementary Information**).

**Figure 5B** (left) presents the results of parameter optimization for different effective dwell times (smart imaging time) and across multiple datasets. This optimization links any effective dwell time (achieved by optimizing the VI for different *T*_initial_, *T*_rescan_) to an accuracy-equivalent standard homogeneous dwell time. For example, an effective dwell time of 150 ns/pixel in the mouse cortex dataset already attains the maximal quality using a specific set of initial, rescan dwell times, and rescan rates that are determined per tile. This quality is comparable to standard homogeneous scan at 800–1000 ns/pixel.

**Figure 5B** (middle) depicts the time saved by SmartEM compared to standard microscopy. For the mouse cortex dataset, the maximal saving compared to standard EM is achieved when SmartEM is used at an effective dwell time of ~125 ns/pixel, which corresponds to an accuracy akin to ~690 ns/pixel by the standard pipeline. This effective dwell time produces images at a speedup of approximately 6× with nearly maximal possible segmentation accuracy (**Figure 1**). The same analysis shows that the *C. elegans* male nerve ring can be acquired at a speedup of approximately 5× and the human temporal lobe at a speedup of approximately 7×.

**Figure 5B** (right) estimates the time to replicate the accuracy of SmartEM using standard microscopy on 1 terapixel of tissue. For the mouse cortex, the SmartEM microscope running for 42 hours of continuous imaging achieves the same quality as a standard pipeline running for 212 hours.

#### Scenario 2: Minimizing imaging time with fixed image quality

In the second scenario, a certain volume needs to be segmented while minimizing imaging cost. The total imaging time is not fixed in advance, but the quality of the SmartEM images must still meet a standard. In practice, SmartEM acquires the volume in a way that achieves segmentation results comparable to standard EM but in significantly less time. First, the operator determines the dwell time required to achieve a specific quality under standard homogeneous scanning, which can be obtained from the SmartEM pipeline’s estimate of a minimum homogeneous dwell time (**Figure 1**). Once the image quality is effectively set by selecting a reference dwell time for uniform scanning, SmartEM then uses its adaptive approach to minimize the overall imaging time while maintaining comparable segmentation accuracy.

We analyzed the expected imaging time of SmartEM across the three datasets by applying the following procedure separately to each tissue. We first acquired images at multiple homogeneous dwell times ranging from 25 to 1200 ns/pixel from the same areas. Next, we applied SmartEM, using the same error detector (ERRNET) and border prediction model (FUSEDEM2B), to produce composite dwell time images derived from different combinations of initial dwell time, rescan dwell time, and rescan rate. To match each standard homogeneous dwell time to an effective SmartEM dwell time, we identified the shortest composite dwell time that produced segmentation results statistically indistinguishable from those of the standard dwell time across tiles (see **Supplementary Information**). **Figure 5C** shows the relationship between the targeted standard dwell time and the SmartEM dwell time with comparable accuracy.

For the mouse cortex, the highest possible quality of standard EM at 800–1000 ns/pixel (see **Figure 1**) is attained by a smart effective dwell time of approximately 149–155 ns/pixel. This ~5.4–6.5× speedup from standard to SmartEM is achieved by selecting the percentage of rescanned pixels in each image tile, and letting ERRNET determine rescan locations. The *C. elegans* male nerve ring, in comparison to standard EM at 800– 1000 ns/pixel, can be acquired with a smart dwell time of approximately 182–207 ns/pixel (~4.4–4.8×). The human temporal lobe, compared to standard EM at 500–1000 ns/pixel, can be acquired at a smart time of approximately 134–154 ns/pixel (~3.7–6.5×).

Image acquisition with widely available SEMs is now a limiting factor in connectomics. This evaluation indicates that the SmartEM pipeline can yield up to ~7× speedup compared to standard image acquisition with an SEM without compromising quality. At competitive short dwell times (~75ns-200ns), SmartEM offers better quality compared to standard EM imaging.

### Imaging and reconstruction of mouse cortex with SmartEM

We applied SmartEM to densely reconstruct multiple portions of mouse cortex tissue. Two volumes of sizes 68×60×3 *µ*m^3^ (**Figure 6A**) and 118×102×3 *µ*m^3^ (**Figure 7**), and a section of size 205×180 *µ*m^2^ (**Figure 6B–J**), were imaged at 4 nm pixel resolution.

**Figure 6.**
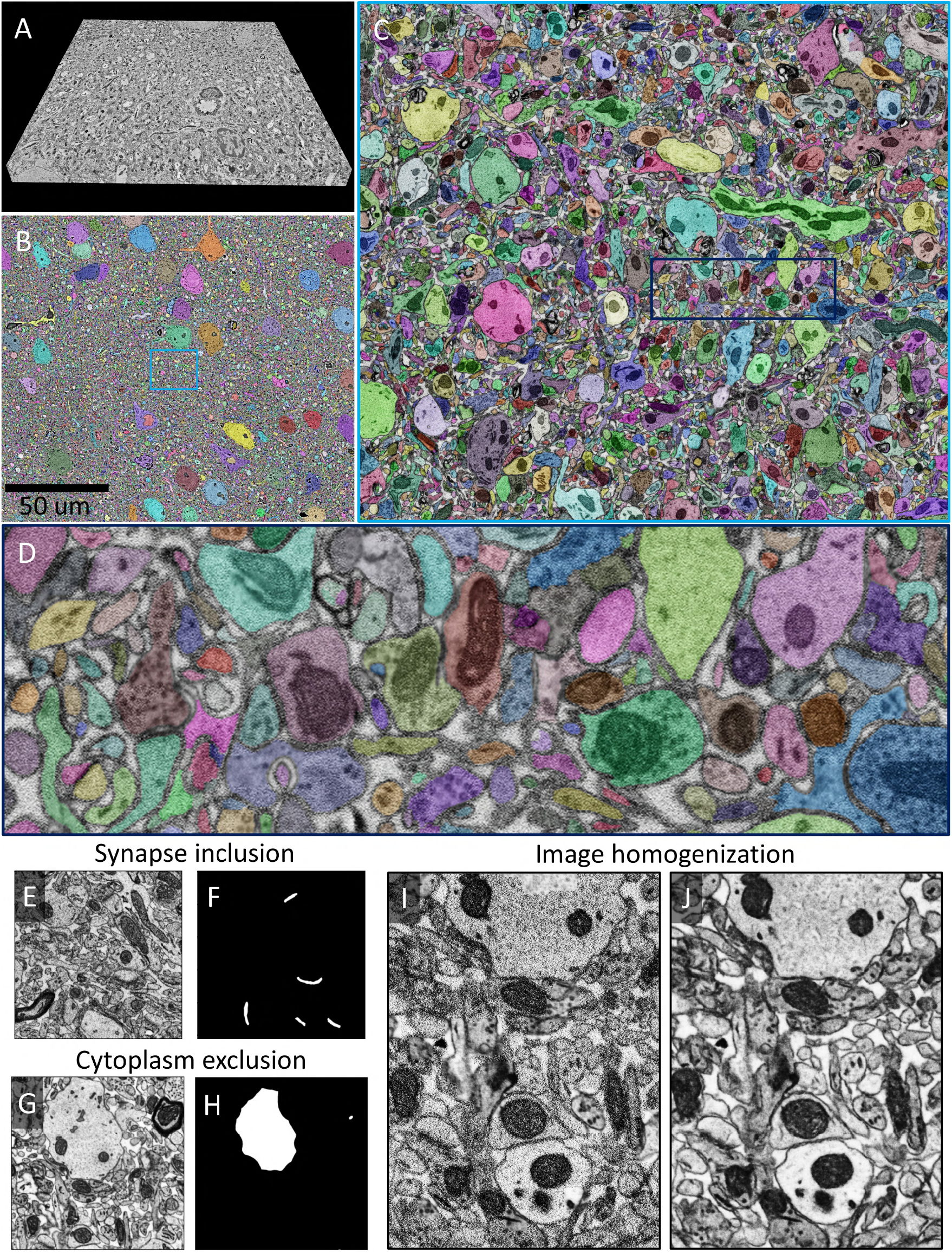
Segmentation of mouse cortex using SmartEM. **A**. Stitched and aligned SmartEM volume of size 68×60×3 *µ*m^3^ ([neuroglancer](https://neuroglancer.bossdb.io/#!https%3A%2F%2Fbossdb-neuvue-datalake.s3.amazonaws.com/public/states/dc6319d0-96db-407a-9e42-05f995567c67.json)). **B**. Segmentation of stitched SmartEM section of size 205×180 *µ*m^2^ using FUSEDEM2B and watershed transform (left panel in [neuroglancer](https://neuroglancer.bossdb.io/#!https%3A%2F%2Fbossdb-neuvue-datalake.s3.amazonaws.com/public/states/dc6319d0-96db-407a-9e42-05f995567c67.json)). **C**. Location of the highlighted region in B with respect to the total section. **D**. Detailed depiction of segmentation in the boxed region in C. **E**,**F**. Automatic detection of synapses from short dwell time images. **G**,**H**. Automatic detection of regions to be excluded from short dwell time images. **I**,**J**. An image (I) made of composite dwell times is stylized to appear akin to a homogeneous dwell time image (J). A comparison between composite dwell time and homogenized images is available in [neuroglancer](https://neuroglancer.bossdb.io/#!https%3A%2F%2Fbossdb-neuvue-datalake.s3.amazonaws.com/public/states/dc6319d0-96db-407a-9e42-05f995567c67.json).

**Figure 7.**
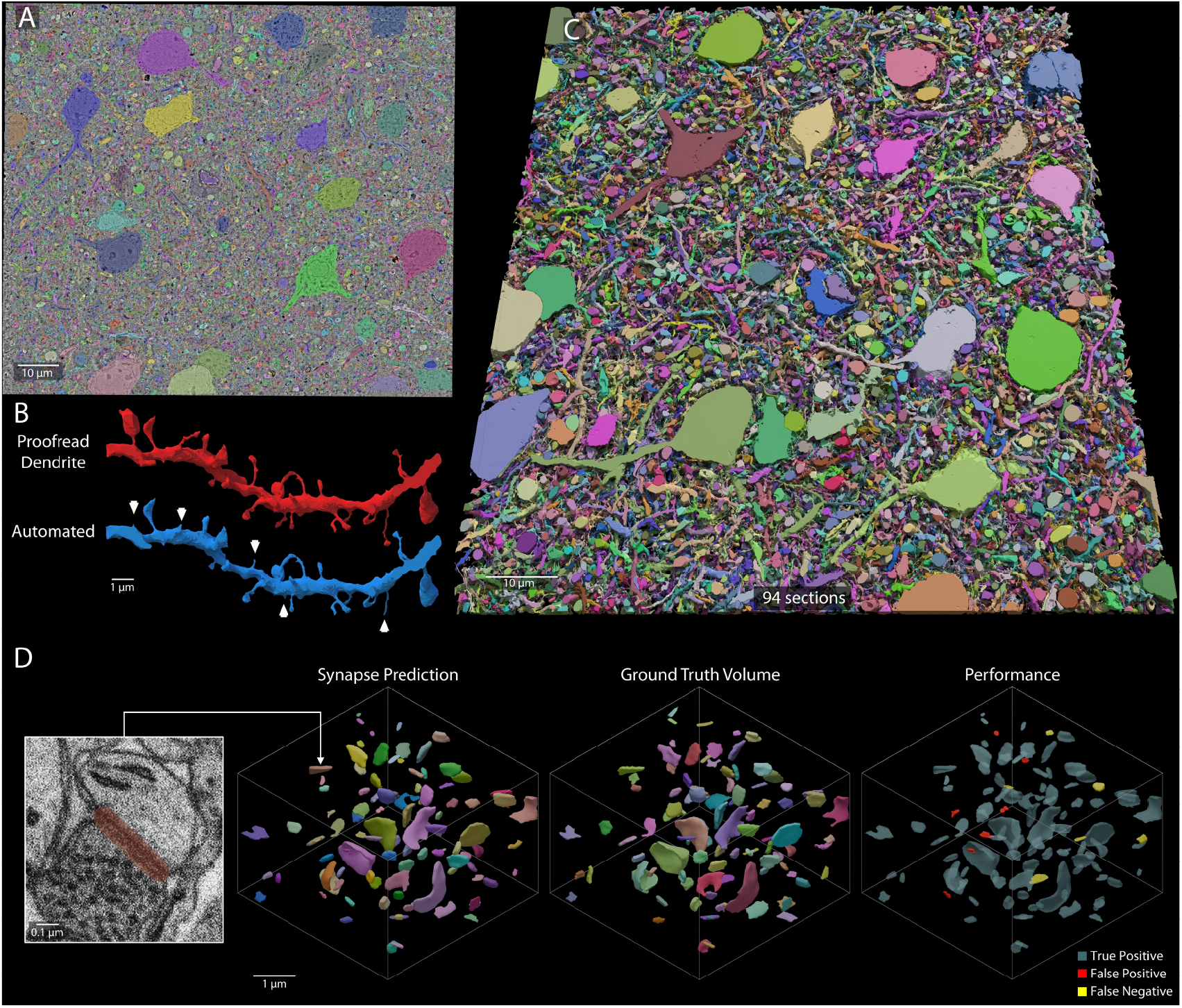
Neuronal and synapse reconstruction of a mouse cortex SmartEM volume taken at an average time of 99 ns/pixel. **A**. A section showing overlay of fused EM and an automated neuronal reconstruction, long and short dwell time pixels at 800 ns/pixel (*T*_rescan_) and 75 ns/pixel (*T*_initial_), respectively, and a rescan rate of 3% (*α*). **B**. A dendrite reconstruction proofread by an expert (red) achieved by manually itemizing and reconstructing all dendritic spines from the fused EM image stack. An automated reconstruction (blue) achieves a high reconstruction rate of the dendritic spines. Arrowheads indicate split errors. **C**. A rendering of the automated 3D reconstruction of all sections in the dataset. The high quality of automated reconstruction has sparse merge errors common to current segmentation algorithms. The reconstruction can be viewed in [neuroglancer](https://neuroglancer.bossdb.io/#!https%3A%2F%2Fbossdb-neuvue-datalake.s3.amazonaws.com/public/states/dc6319d0-96db-407a-9e42-05f995567c67.json). **D**. Smoothed renderings of synapses showing the neural network prediction, expert ground truth and a comparison. In the comparison, true positives are labeled in opaque light blue, false positives in red and false negatives in yellow. Synapse network predictions and ground truth can be visualized in [neuroglancer](https://neuroglancer.bossdb.io/#!https%3A%2F%2Fbossdb-neuvue-datalake.s3.amazonaws.com/public/states/dc6319d0-96db-407a-9e42-05f995567c67.json).

For the first volume acquisition, we used an initial dwell time of 75 ns/pixel, rescan dwell time of 800 ns/pixel, and rescan rate of 10%, providing an effective dwell time of

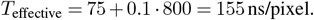

This average dwell time for SmartEM corresponds to a standard dwell time of ~1000 ns/pixel for traditional microscopy (see previous section). This acquisition tested the ability to acquire, stitch and align in 3D serial-section volumes.

For the second volume acquisition, we employed even more competitive SmartEM parameters with an initial dwell time of 75 ns/pixel, rescan of 800 ns/pixel, and a rescan rate of 3%, providing an effective dwell time of *T*_effective_ = 75 + 0.03 *·* 800 = 99 ns/pixel.

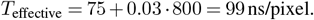

For this volume, a comparison between the co-registered EM images of short dwell time and composite dwell time is available in [neuroglancer](https://neuroglancer.bossdb.io/#!https%3A%2F%2Fbossdb-neuvue-datalake.s3.amazonaws.com/public/states/dc6319d0-96db-407a-9e42-05f995567c67.json). This acquisition tested whether highly competitive SmartEM imaging parameters would support accurate automated neuronal reconstruction in 3D (described below).

To test the scalability of SmartEM to larger imaging grids, we acquired a section of size 205 × 180 *µ*m^2^ composed of 30 × 30 individual tiles with an initial dwell time of 75 ns/pixel and a rescan of 600 ns/pixel and a rescan rate of 10%, providing an effective dwell time of

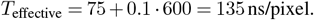

As mentioned above, this effective dwell time corresponds to the maximal possible speed up of SmartEM for this dataset, producing images with segmentation quality akin to standard EM at ~1000 ns/pixel. We depict the segmentation of pipeline outputs in **Figure 6B, 6C, 6D** (left panel in [neuroglancer](https://neuroglancer.bossdb.io/#!https%3A%2F%2Fbossdb-neuvue-datalake.s3.amazonaws.com/public/states/dc6319d0-96db-407a-9e42-05f995567c67.json)).

We also assessed the ability to detect synapses on short dwell time images (from 25 to 1000 ns/pixel) and applied this detection to the above initial scan of 75 ns/pixel with excellent results that are comparable to long scan imaging as shown in **Figure 6E, 6F, S6**. In **Figure 6G, 6H**, we show the ability of SmartEM to detect and exclude regions of no interest, where cytoplasm far from membrane is detected from the initial scan, allowing SmartEM to force the skipping of the long dwell time scanning from these regions. In **Figure 6I, 6J, S9**, we demonstrate the ability to translate the fused images to uniform looking EM tiles with quality akin to long dwell time imaging (visualized in [neuroglancer](https://neuroglancer.bossdb.io/#!https%3A%2F%2Fbossdb-neuvue-datalake.s3.amazonaws.com/public/states/dc6319d0-96db-407a-9e42-05f995567c67.json)).

### Neuronal reconstruction of mouse cortex

We tested SmartEM in application to connectomics. Connectomics requires accurate agglomeration of 2D cross-sections (see **Figure 7A**) into 3D reconstruction of neuron volumes and synapses (see **Figure 7C, 7D**). We asked whether the quality of aligned SmartEM fused images supports automated reconstruction and proofreading with comparable performance to traditional imaging. We first focus our analysis to the problems of neuron reconstruction. We applied a lightweight 3D neuron segmentation algorithm (see **Supplementary Information: Segmentation and neuronal reconstruction**) to the mouse cortex volume acquired at a competitive average dwell time of 99 ns/pixel (visualized in [neuroglancer](https://neuroglancer.bossdb.io/#!https%3A%2F%2Fbossdb-neuvue-datalake.s3.amazonaws.com/public/states/dc6319d0-96db-407a-9e42-05f995567c67.json)). We assessed the quality of resulting SmartEM image volume with automated reconstruction of fine processes and expert manual annotation (**Figure 7B**), as described below.

### Reconstruction of dendritic spines

Connectomes can contain “split” errors (fragmenting the volume of one cell) or “merge” errors (joining the volume of two cells). Because a comprehensive analysis of merge errors typically requires larger reconstructed volumes to assess metrics such as error-free runlength, we qualitatively inspected and verified that none of the large segmented objects was implicated in catastrophic merge errors (see **Figure 7C**). Spines are the fine processes that protrude from dendrites and contain synapses. To further benchmark SmartEM performance quantitatively, we studied split errors in the 3D reconstruction of dendritic spines, a challenging feature for automated reconstruction. We randomly selected three dendrites (see **Figure 7B**). We counted spines that were fully automatically reconstructed without split errors and spines with split errors. Expert human annotators verified every correct reconstruction and verified that every split error was correctable with proofreading. The percentage of correct spines was approximately 58%, 53% and 75% in the three dendrites. The combined percentage of correct spines was 65%, comparable to the rate of correct spine capture in recent automated reconstruction of human cortex (67%) (Shapson-Coe et al., 2024).

### Reconstruction of synapses

In addition to validating the quality of automated neuron reconstruction in the mouse cortex volume, as described above, we also trained a neural network to automatically reconstructed synapses (see **Supplementary Information: Synapse segmentation technique**) and validated the results against expert manual annotation (**Figure 7D**, [neuroglancer](https://neuroglancer.bossdb.io/#!https%3A%2F%2Fbossdb-neuvue-datalake.s3.amazonaws.com/public/states/dc6319d0-96db-407a-9e42-05f995567c67.json)). We measured object-wise synapse precision, recall and the F1-score in 3D (see **Supplementary Information: Synapse validation**). When evaluated on the test dataset, we obtained a precision of 93.2%, a recall of 94.1% and an F1-score of 93.7%, comparable to state-of-the-art performance on traditional EM volumes (Turner et al., 2020; Bae et al., 2025).

## Discussion

Recent advances in machine learning and cloud computing are shifting the bottleneck in connectomics from image analysis to data acquisition. While cutting-edge connectomics now generates datasets at cubic millimeter scale and beyond, progress toward reconstructing large-scale brain tissues or democratizing connectomics across neuroscience labs remains hampered by imaging throughput. This acquisition bottleneck is likely to become even more pronounced as ambitions expand toward larger connectomes produced in greater numbers.

The SmartEM approach described here directly addresses this challenge by integrating computational intelligence into single-beam scanning electron microscopes. Entirely implemented using commodity computer hardware, SmartEM has the potential to transform widely available, affordable single-beam SEMs into high-throughput connectomic platforms with minimal hardware modification. In our experiments, SmartEM increased imaging throughput by up to ~7-fold compared to traditional SEM imaging methods, without loss in segmentation accuracy.

Beyond accelerating imaging acquisition, SmartEM’s computational framework is adaptable to different microscopy modalities, enabling intelligent, data-aware imaging in various scientific fields (see below).

### The flexibility of SmartEM

The SmartEM pipeline can be flexibly reconfigured to meet the needs of different users and diverse sample preparations. First, SmartEM allows an SEM to identify error-prone regions in any rapidly acquired image using the ERRNET neural network. A key feature of ERRNET is that it can be trained using any independently defined segmentation algorithm to detect discrepancies between rapidly acquired and high-quality, slowly acquired EM images. Our implementation uses variation of information to quantify segmentation differences, but other error detection schemes and segmentation methods can be substituted during training, depending on laboratory preference. Second, SmartEM allows an SEM to perform a long dwell time rescan of any region within an initially rapidly acquired image in real-time during microscope operation. This rescan can be done with any SEM equipped with electrostatic scan generators capable of deflecting the electron beam to any pixel in an image much faster than the shortest dwell time per pixel (e.g., <25 ns/pixel) (Mohammed and Abdullah, 2018; Anderson et al., 2013). Third, SmartEM can segment multi-dwell time images using the FUSEDEM2B neural network. The training pipeline for this network is adaptable to other segmentation algorithms.

### Diverse use cases for SmartEM

The underlying concept of SmartEM can improve the efficiency and accuracy of SEM image acquisition in any context where it is beneficial to selectively adjust imaging time across different regions. Much like the human eye, which captures most of a visual scene with rapid low-resolution (non-foveal) imaging and selectively dwells with high-resolution (foveal) imaging on specific areas to resolve ambiguity (Thorpe et al., 1996), SmartEM rapidly scans entire regions and selectively rescans areas predicted to require higher fidelity. SEM is widely used in materials science and manufacturing, where samples also often have regions varying significantly in detail and complexity. These applications, as well as others where specific structural features can be predicted but not accurately reconstructed from an initial rapid scan, are especially suited to SmartEM. Imaging approaches that take advantage of electron beam-sensitive materials, such as cryo-EM, could particularly benefit from the selective rescanning of SmartEM. Sparsely distributed structures or molecules of interest can first be rapidly identified and then selectively rescanned at longer dwell time, reducing overall beam exposure while enhancing image quality.

Although we focused here on neuronal reconstruction for connectomics, SmartEM was also adapted to selectively rescan high-quality images of salient structures like chemical synapses, providing morphological reconstructions without significant increases in total imaging time. Likewise, SmartEM can be readily adapted for applications in cell biology or pathology by selectively recognizing and rescanning other sparse but biologically important structures, such as mitochondria or other organelles.

The SmartEM pipeline can not only be “taught” to capture the most salient features of an image, but can also be used to neglect regions without interest. In most connectomics of larger brains, nearly all objects in the field of view will be neural structures. But in small invertebrates, neural tissue might constitute only a small part of the field of view. The *C. elegans* nerve ring (brain) is <10% of the total volume of the body, and wraps around the pharynx. Any two-dimensional brain section of the *C. elegans* nervous system will also include substantial nonneural tissue. To date, connectomic datasets have been acquired by carefully designating the region-of-interest for each image. In such cases, SmartEM could significantly simplify and accelerate image acquisition by automatically focusing scanning time on neuronal tissue, eliminating the laborious manual specification of regions of interest by users.

### Adaptability of SmartEM for other microscopes and other applications

Tape-based serial-section sample collection, where specimens are stored permanently and can be re-imaged at any time, is suited to SmartEM because any information that is lost during imaging can be recovered. When specimens are imaged for the purpose of connectomics, the SmartEM pipeline might gloss over features that might eventually be of interest to other scientists for other applications (e.g., cell biology). Because serial-sections stored on tape can be safely archived for years, they can be revisited at any time.

Instead of collecting serial sections on tape, one can use block face imaging with serial tissue removal. One block face approach, Focused Ion Beam SEM (FIB-SEM), has distinct advantages over tape-based serial-section sample collection, including thinner tissue layers (4-8 nm) and better preservation of image alignment (Knott et al., 2008). The principal disadvantage of FIB-SEM has been the slow pace of traditional SEM with >1000 ns/pixel dwell times. This can be problematic when the microscope is used to collect extremely large specimens, and must be continuously operational for days or weeks without technical glitch. However, a FIB-SEM that implements the SmartEM pipeline would be able to operate much faster, increasing the likelihood of capturing an entire specimen in single long runs. SmartEM is expected to provide greater speedup on block face imaging because the imaging component is a larger part of the entire acquisition pipeline compared to serial-section SEM. Similar benefits will be obtained with other block face imaging approaches such as Serial Block Face SEM (SBF-SEM) where a diamond knife slices the specimen (Denk and Horstmann, 2004).

### Improvements for SmartEM

Advances in segmentation algorithms and microscope hardware are expected to enable further reduction, potentially by an order of magnitude, in total beam time with SmartEM, without compromising segmentation accuracy. Such gains may come from significantly shorter dwell times, below current minimums of 25 ns per 4 nm pixels. In addition, the need for rescans may be lessened by leveraging 3D contextual information. To do so, SmartEM can (1) use the 3D context from adjacent sections to remove redundancies in rescan masks of adjacent sections, such as by detecting aberrant neuronal morphologies (Rolnick et al., 2017; Zung et al., 2017; Celii et al., 2025), and (2) image most of the neuropil at 8 nm pixels, while scanning and segmenting large neuronal elements including cell bodies at a much lower resolution (e.g., >16 nm) (Shapson-Coe et al., 2024). Intelligently adjusting spatial resolution with SmartEM, however, would require careful investigation of how different beam currents influence electron beam spot size and, consequently, segmentation accuracy. We note that commercial multibeam SEMs, with their multiple beams controlled synchronously, cannot directly leverage some of these SmartEM strategies. Nonetheless, our innovations could significantly accelerate single-beam SEMs, positioning them as a viable alternative to the currently used high-throughput electron microscopes for connectomics.

## Code Availability

Machine learning software and all models will be made available upon publication on a public repository. and are currently available on request.

## Data Availability

SmartEM data is released on BossDB (Meirovitch et al. Project Page for details). Additional information on this BossDB data collection is provided in the data accessibility supplementary section.

## Acknowledgements

Research reported in this paper was supported by the NIH BRAIN Initiative under award number 1U01NS132158 and by NIH grants 5U24NS109102 and U01 NS108637. L.M.’s work was supported in part by a fellowship from MathWorks. T.A. is supported by the MIT-Novo Nordisk Artificial Intelligence Postdoctoral Fellows Program.

## Supplementary Information

### Segmenting composite images

The smart microscope should be able to analyze images composed from multiple dwell times (see **Figures 1C, 2B, 2C, 3, 6A-6D**). We tested whether replacing error-prone regions in a short dwell time image with regions taken from long dwell time images improves segmentation outcomes. **Figure S1** depicts the segmentation outcome of a short dwell time image taken at 100 ns/pixel segmented with a dedicated 100 ns/pixel network FASTEM2B (**S1A,S1E**), and by FUSEDEM2B (**S1B,S1F**).

The segmentation quality of these networks are similar (top panel; VI=0.025 and VI=0.022). In most scenarios, the network trained to deal with fused EM (FUSEDEM2B) produces better results than networks trained to handle a fixed dwell time, even if the input to the two networks consists of a single homogeneous dwell time. **Figures S1C, S1G** depict the segmentation of an image where the error-prone regions were detected by an error detector and replaced with long dwell time pixels (2500 ns). The error level is typically and substantially cut by ~3-4×. The 2500 ns/pixel reference image and its segmentation are shown in **Figures S1D, S1H**. All error estimates based on VI shown in **Figure S1** are presented as the sum of the merge error term and split error term.

### Imaging procedure

The SEM is automated to acquire images of individual tiles of every specimen section that are eventually stitched and aligned to form a total image volume (**Figure 3**). The microscope navigates through multiple specimen sections held on tape and defines every specimen region of interest (S-ROI). Each S-ROI is captured at high spatial resolution by multi-tile acquisition. To identify the S-ROI and automate stage position and rotation control, we used SEM Navigator, a custom interface akin to earlier WaferMapper software (Hayworth et al., 2014). The list of S-ROIs is exported into a text file, which is subsequently processed by the SmartEM pipeline (coded in Python/Matlab) using the Thermo Fisher Scientific Autoscript (Thermo Fisher Scientific, 2018) package. The SmartEM pipeline controls the Verios (Thermo Fisher Scientific, 2020) microscope, moves to S-ROI and individual tile positions, controlling the entire acquisition sequence.

For all image acquisitions, we used the Verios UHR (Ultra High Resolution) imaging mode with 4 nm/pixel spatial resolution and ~ 4 mm working distance. Image contrast was obtained using a back-scattered electron detector with 2000 V stage bias. The initial short dwell time scan was obtained using the full frame acquisition Autoscript interface. The subsequent long dwell time rescan utilized the standard interface of Autoscript patterning To optimize image quality and tuning time for both short movements between neighboring tiles and long movements neighboring sections, we customized sequences of various autofunctions. These autofunctions included autocontrast/brightness (ACB), auto-focus (AF), auto-stigmation (AS), auto-focus/stigmation (AFS), and auto-lens (AL) alignment.

Because we used different interfaces for the initial short dwell time scan and long dwell time rescan, an additional alignment procedure was necessary to achieve pixel-resolution precision in the rescan. The basic system configuration for the rescan acquisition is described in Potocek (2021).

When the rescan long dwell time was shorter than ~ 500 ns/pixel, an unavoidable artifact due to limited system response of the electron deflection system occurred at the edge of rescan regions. We excised this artifact by omitting a 1-pixel boundary from every rescan region.

### Segmentation quality metric

To compare the segmentation quality of different samples we used a variation of information (VI) metric (Meila, 2003). In principle all comparisons that we made in this study can be accomplished with other metrics of segmentation quality as long as they can be applied to 2-dimensional images. We expect the choice of segmentation metric to have little effect as long as any metric assesses similar topological attributes as VI (i.e., whether objects are split or merged). Our implementation of the VI running on CPU/GPU is available at https://pypi.org/project/python-voi/.

### Using VI to build ERRNET

To train the error detectors we needed to locate the specific regions that contribute to the largest segmentation differences between image pairs, which is not provided by the VI metric. VI combines split and merge errors. The two error measures are defined by comparing the entropy of three segmented images (Meila, 2003), 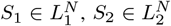 and 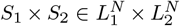 for two *N*-pixel labeling (instance segmentation) *S*_1_ and *S*_2_ that needs to be compared, where the *L*s represents the sets of pixel labels. The segmented image *S*_1_ × *S*_2_ is labeled by concatenating the labels from *S*_1_ and *S*_2_ for each pixel. The VI is then the sum of two error terms VI_merge_ and VI_split_ VI = VI_merge_ + VI_split_.

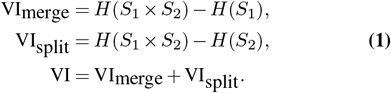

Due to the additivity of the entropy measure (Meila, 2003), VI_merge_ and VI_split_ can be broken into individual constituents, representing the amount of error contributed by each individual label in each segmentation. We could thus rank objects in each segmentation according to the amount of variation they contribute to overall VI (**Figures S2**). The error contributed by the set of pixels that are both in segment *s*_1_ *∈ S*_1_ and *s*_2_ *∈ S*_2_ (i.e. the error contributed by a segment in *S*_1_×*S*_2_) is

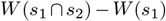

and

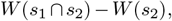

for the split and merge errors, respectively, where 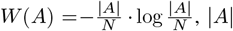, *A* is the number of pixels in *A* and *N* is the number of pixels in the image.

**Figure S1.**
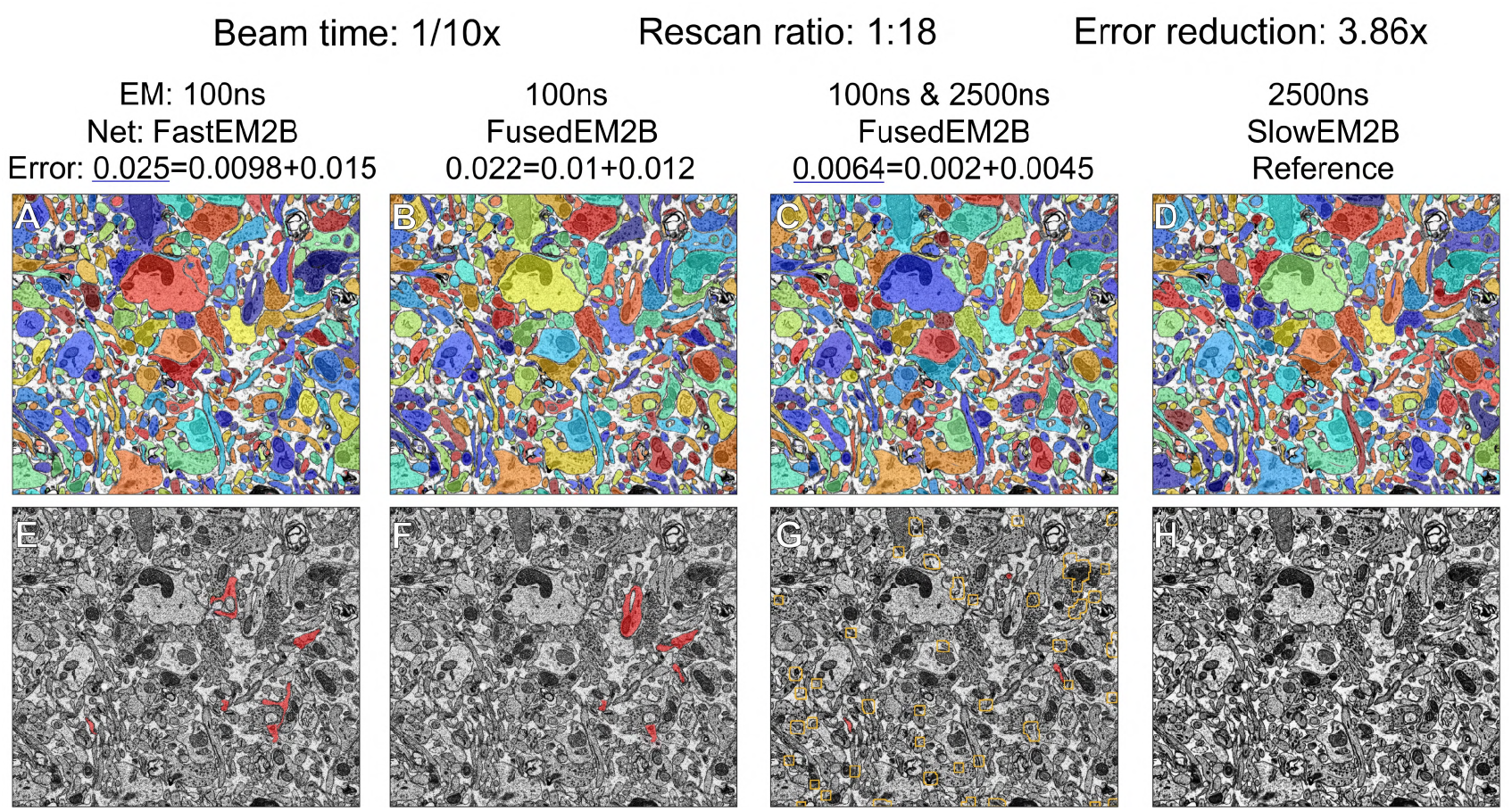
Composite EM images fusing short and a long dwell time regions are better segmented compared to short dwell time images. We tested in the mouse cortex datasets whether replacing error-prone regions harms the ability to segment. Composite images tend to be segmented with significantly higher accuracy. Error of the instance segmentation is assessed in terms of the Variation of Information (VI) compared to the segmented reference image, where VI is composed of a merge and split error terms as in Equation 1.

Once the significantly incompatible objects are detected in each segmentation, we used image processing to delineate the borders that are responsible for the topological differences between the two segmented images (**Figure S3**). We then produced binary masks from these errors and trained neural networks (ERRNET) to detect them directly from border probability maps, themselves produced by another neural network (FASTEM2B). Detecting borders allows our technique to disregard small “cosmetic” variations between two segmentations that do not cause meaningful topological differences.

### Determination of maximal segmentation quality

We developed an unbiased estimate for the minimal dwell needed for 2D segmentation. We compared segmentations from *N* images for each pair of dwell times *d*_1_ < *d*_2_ and an overly slow dwell time *d*_ref_. We asked whether the VI of the *d*_2_ images was significantly smaller (p<0.05; Wilcoxon signed rank test) than *d*_1_ images compared to *d*_ref_ images. When two dwell times were not sufficiently different, we call these dwell times equivalent. We defined the minimum dwell time with near maximal segmentation ability as that dwell time beyond which VI does not improve.

### Forcing fast scan imaging of desired regions

The acceleration of SmartEM depends on the quantity of rescanned pixels. Since the rescanning mask is learned rather than calculated through a fixed process, regions irrelevant to the connectomics task may contain error-prone regions and appear in the rescan map, potentially reducing speedup. To exclude irrelevant regions from slow rescan, we built another neural network module (EMEXCLUDE) to calculate what regions should be excluded from any rescan, even if they might be flagged as error-prone by ERRNET. Developing a separate EMEXCLUDE module (rather than adding this capability to ERRNET) conferred additional flexibility to the SmartEM pipeline by allowing us to adaptively choose what regions should be excluded from rescan without retraining ERRNET. Bypassing irrelevant pixels (e.g., cell nuclei, blood vessels) during rescan boosts efficiency by conserving time and computational resources.

Here, we implement EMEXCLUDE to exclude regions that are sufficiently far from any cellular borders. To do this, we utilize the Euclidean distance transform on input binary borders. This transform calculates the shortest Euclidean distance from each zero pixel (background) to any non-zero (foreground) pixel in the image. To train EMEXCLUDE, we binarize the distance transform with a fixed threshold (**Figure S4**). The features of irrelevant regions we learned as a semantic segmentation task using paired EM images and their binary masks (see **Neural network models**). The SmartEM pipeline applies EMEXCLUDE in real-time on short dwell time images and precludes rescanning irrelevant regions that might have been predicted by ERRNET. To assess the performance of different modules in the SmartEM pipeline, we exclude EMEXCLUDE from speedup tests shown in **Figure 5**. For the cytoplasm exclusion described above, the average exclusion proportion is about 23% as shown in **Figure S5**. The speedup tests shown in **Figure 5** would improve with the implementation of EMEXCLUDE.

**Figure S2.**
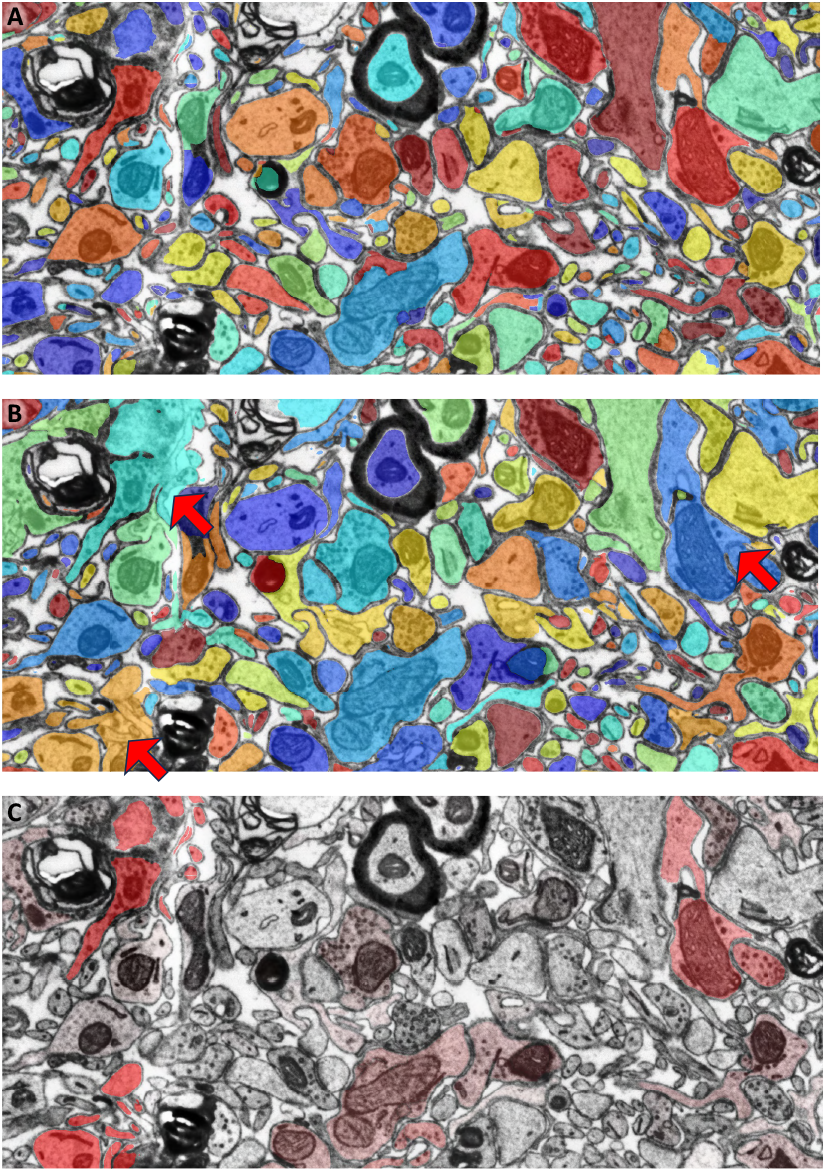
Ranking objects of two segmented images based on contribution to variation of information. **A**. Segmentation of long dwell time image at 1000 ns. **B**. Segmentation of short dwell time image at ~ 100 ns/pixel overlaid on 1000 ns/pixel EM. Some large errors are indicated with red arrows. **C**. Objects that vary between the two segmented images. Red heatmap indicates contribution to variation of information (Meila, 2003) where variable objects come from either of the two segmented images. The largest variation is captured by the three objects indicated by red arrows.

### Identifying additional high-interest regions for slow rescan

ERRNET identifies regions susceptible to segmentation errors and rescans them at a higher quality to improve segmentation accuracy. The same strategy can be re-formulated, not only to identify error-prone regions, but to identify additional image-specific regions of special interest, such as synapses or any sub-cellular component of biological interest. Here, we built an additional neural network module (EMINCLUDE) to rescan regions identified as synapses, because of their high relevance to connectomics. Mouse cortex typically contains ~1-1.5 synapses per *µ*m^3^ (Kasthuri et al., 2015), or ~2-3 synapses per field of view when image tiles are ~8 × 8 *µ*m^2^. Because of synapse sparsity, the rescan time does not substantially increase. We trained EMINCLUDE with a set of manually-annotated long dwell time SEM images.

To train EMINCLUDE, we first trained a neural network to detect synapses using manual annotations of long dwell time images (EMINCLUDE). The high performance of EMINCLUDE is shown in **Figure S6**. We paired short dwell time images with the binary masks for synapse locations predicted by EMINCLUDE (which had used long dwell time images to make the predictions). This procedure created ground truth to train EMINCLUDE. A snapshot of the synapse detection and rescan mask generation pipeline is shown in **Figures S7,S8**. The hyper-parameters and training details of EMINCLUDE are similar to EMEXCLUDE.

**Figure S3.**
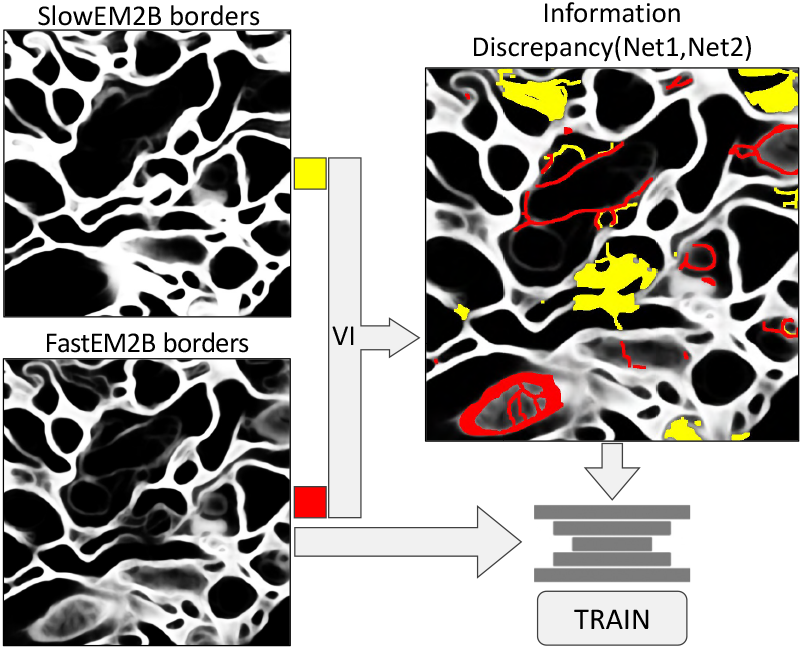
The discrepancy between segmentation with long dwell time (using SLOWEM2B) and short dwell time (using FASTEM2B) is defined based on VI. VI is the sum of individual error terms contributed by each object in the two segmented images. The most variable objects are flagged. Image processing is used to delineate specific borders that appear in only one segmented image. Yellow represents segmented objects that are uniquely predicted in the long dwell time image. Red represents segmented objects that are uniquely predicted in the short dwell time image. A neural network (ERRNET) is trained to predict all red and yellow discrepancies only using short dwell time images. This is possible because variation occurs where border predictors are uncertain and often with typical, at times biologically implausible, border prediction.

**Figure S4.**
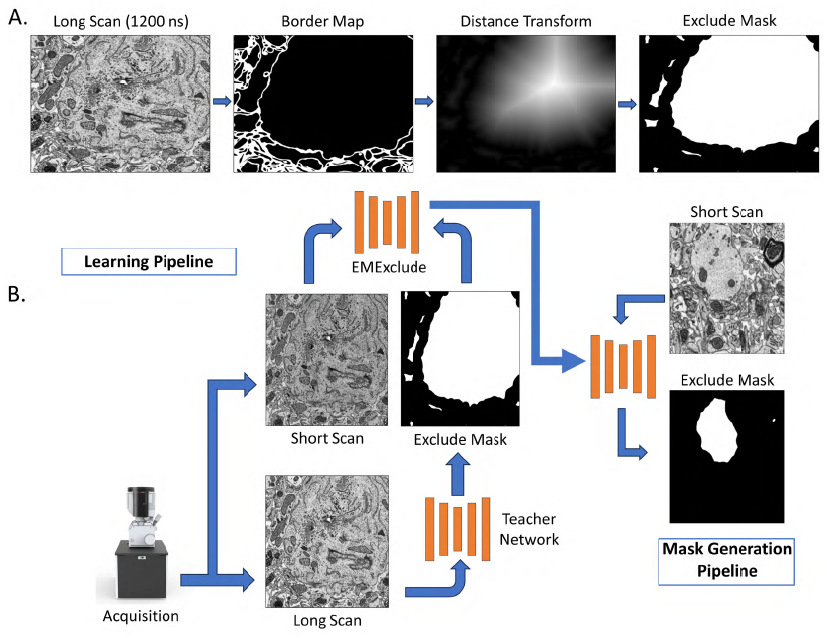
A. The process of generating the EMEXCLUDE ground truth: EM images taken at long dwell times are processed to determine regions that should be excluded in subsequent scans. The sequence begins with the raw EM image, proceeds to border predictions highlighting essential structures, and then applies a Euclidean distance transform to emphasize key features. The final output is a binary differentiation after thresholding, which identifies areas of minimal interest, establishing the EMEXCLUDE ground truth. B. The EMEXCLUDE ground truth is paired with fast EM images to train a neural network, enabling it to recognize and exclude similar non-essential regions in new scans. Once trained, the network processes new EM images in real-time, generating EMEXCLUDE masks.

**Figure S5.**
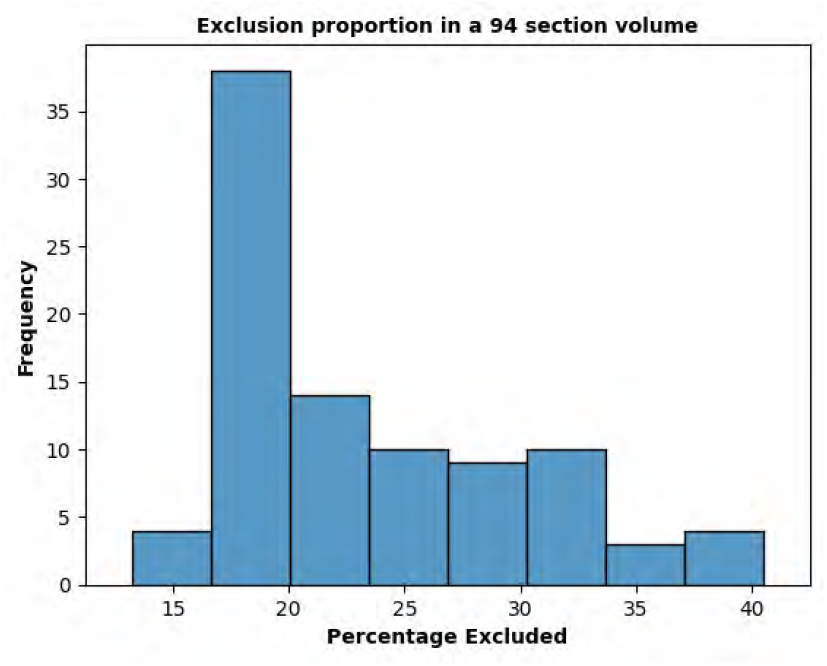
Percentage of EM that can be excluded in a section of the mouse cortex dataset (volume 68 × 60 × 3µ*m*^3^). On average, around 23% of the volume can be excluded from rescanning.

**Figure S6.**
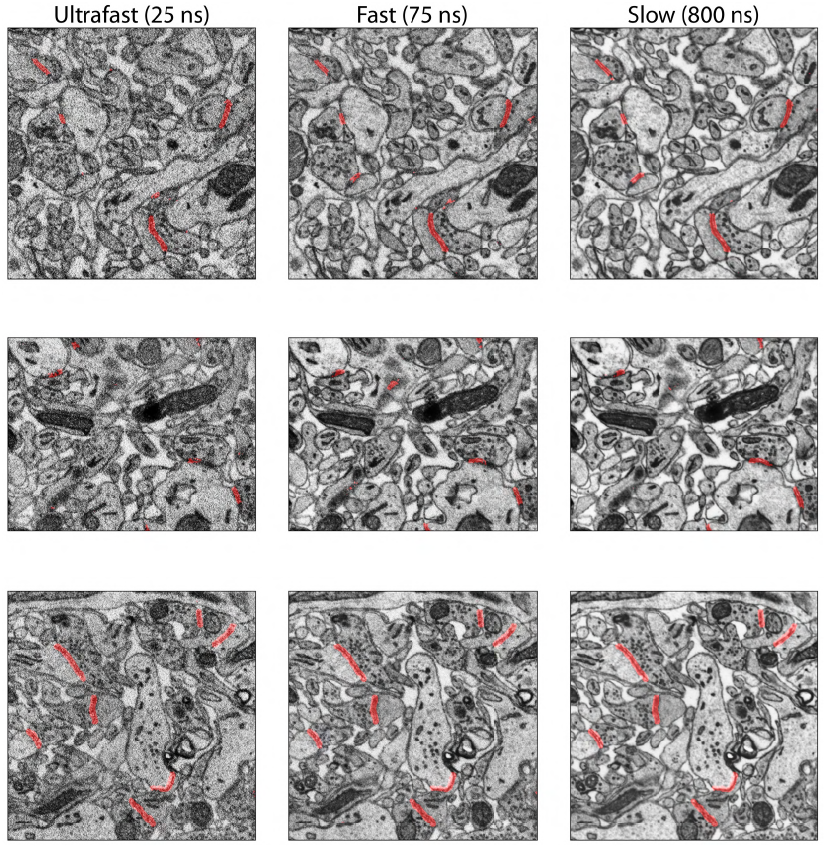
Synapse detection in ultrafast (25 ns), fast (75 ns) and slow (800 ns) dwell time. EMINCLUDE works at multiple dwell times.

### Optional image homogenization

The SmartEM pipeline produces composite image with pixels acquired at different dwell times. A human observer will note contrast differences at interfaces between pixels with different dwell times. To increase human image interpretability, we built an image translator component that homogenizes SmartEM images to look like standard EM images with uniform dwell times. **Figure S9** shows a specific example, a fused EM image that is a mosaic of sub-images with different dwell times. To mitigate dwell time contrasts and produce a visually coherent image, we applied a conditional generative adversarial network (IMAGE-HOMOGENIZER, cGANs) (Mirza and Osindero, 2014). Previous studies used deep learning to improve the quality of microscopy images (Fang et al., 2021; Wang et al., 2019; Weigert et al., 2018; Mi et al., 2021), de-noise EM images (Minnen et al., 2021), and perform image reconstruction across different modalities (Li et al., 2023). IMAGEHOMOGENIZER contains two convolutional neural networks (CNN): a *generator* and a *discriminator* (Isola et al., 2016). Training data are a composite image and a uniformly long dwell time image, where the composite image is generated by randomly combining pixels from short dwell time and long dwell time images in different proportions (**Figures 6B,6C,6D** where the composite images consist of 75 and 600 ns/pixel dwell times). As shown in **Figure S10**, during the training process, the *generator* translates the simulated composite images to resemble long dwell time images, and the *discriminator* attempts to distinguish the translated images from real long dwell time images. The training process utilizes L1 loss and adversarial loss. After image homogenization by the *generator*, the fused EM images are more suitable for human inspection and retain the visual details of fine ultrastructure **Figure S9**.

**Figure S7.**
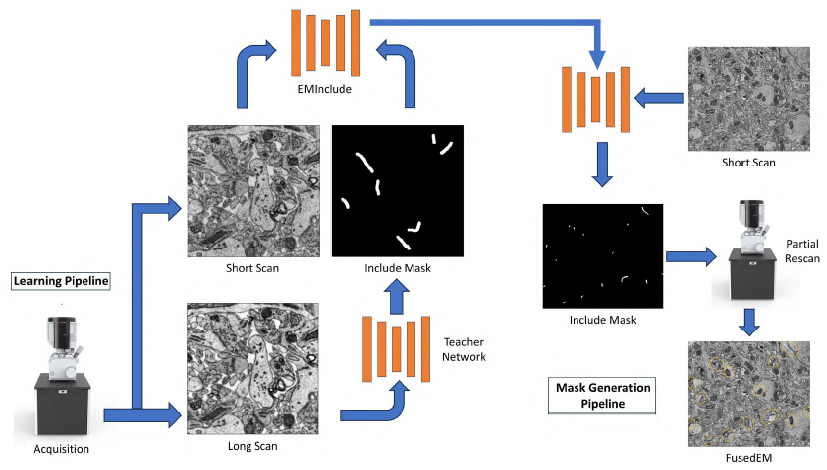
Synapse detection and rescan mask generation pipeline: aligned acquisition provides electron microscopy (EM) images at varying dwell times. A teacher network is trained to identify synapses from slow dwell time images, and these identified labels train a student network, EMInclude, for synapse detection on faster dwell time images. This student network predicts synapse locations to generate a rescan mask, directing the microscope for targeted slow point scans of selected synapses. The outcome is a fused EM image that integrates different dwell times, optimizing scanning speed and detail in areas of interest.

**Figure S8.**
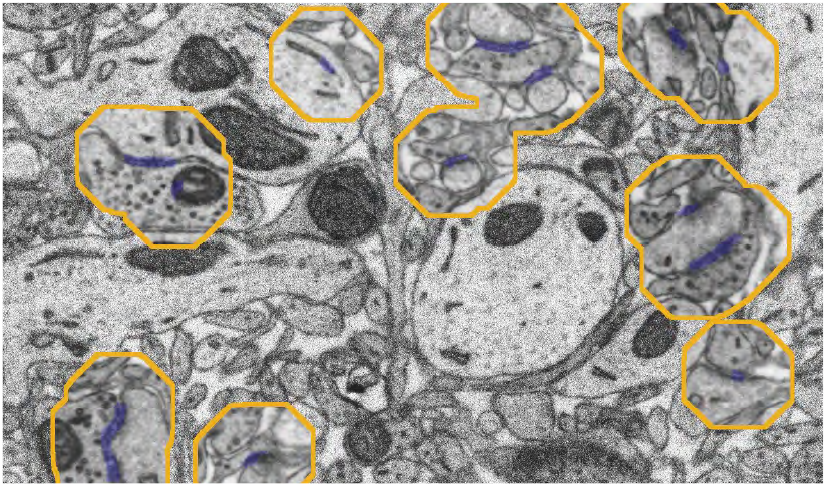
Data-aware imaging of synapses at long dwell time. SmartEM takes a short dwell time image (50 ns/pixel), predicts locations that contain synapses, and rescans these regions at long dwell time (1200 ns/pixel). The blue overlay presents synapse predictions by EMINCLUDE. Yellow outlines represent locations for rescan based on dilation of EMINCLUDE predictions.

**Figure S9.**
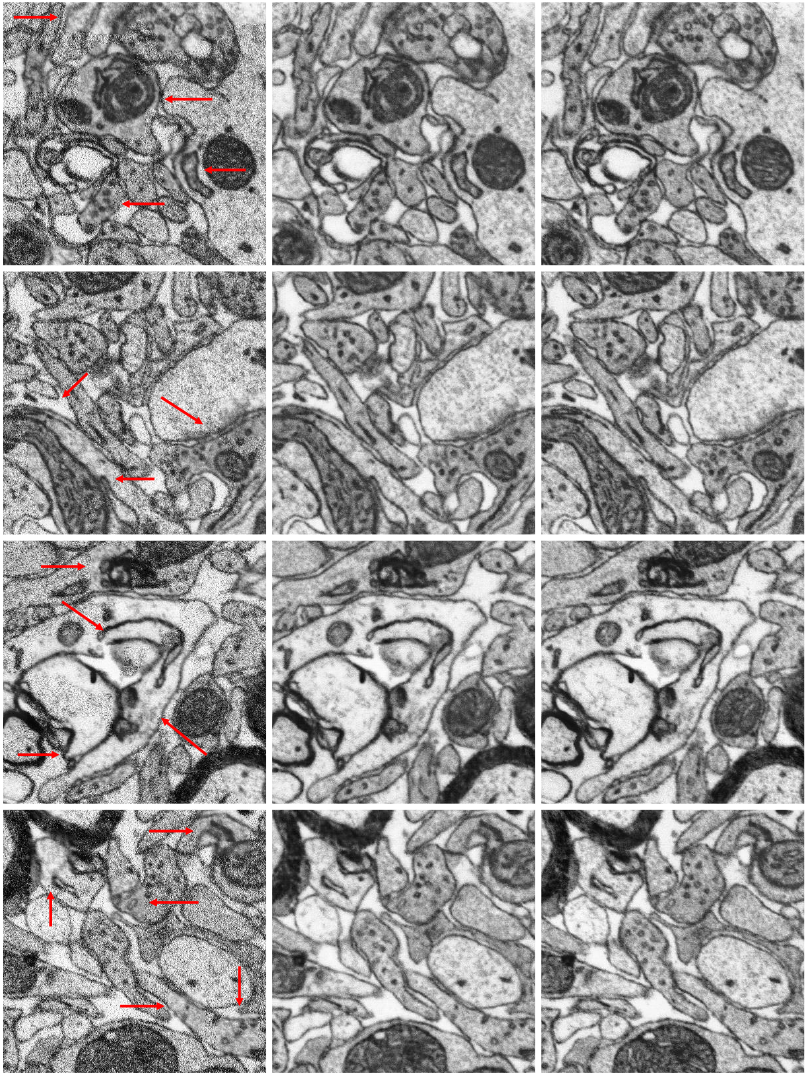
Examples of image homogenization by IMAGEHOMOGENIZER. Left column: composite EM with two dwell times (75 ns/pixel and 600 ns/pixel). Middle column: homogenized EM from composite EM, exhibiting similar visual coherence compared to slow EM. Right column: slow EM (600 ns/pixel). Red arrows indicate the locations with slow dwell time of 600 ns/pixel in composite EM.

**Figure S10.**
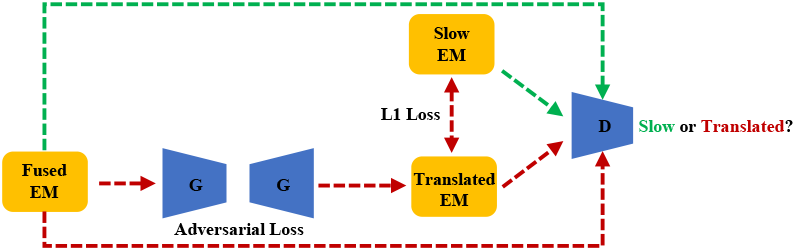
Image Translation Model. G: *generator*. D: *discriminator*. The generator G takes a fused EM as input and produces a translated EM (i.e., fake slow EM) that looks similar to slow EM (i.e., taken by the microscope). The discriminator D takes as input a concatenation of a fused EM and another image that is either slow EM (green arrows) or a translated EM (red arrows). The aim of the discriminator is to classify whether the second image is slow EM or translated EM. The model is trained with a combination of adversarial loss and L1 loss.

### Neural network architectures

For all neural network models, we strove for simple architectures that would allow straightforward reproducibility of results. A U-Net like architecture (Ronneberger et al., 2015) was used to train border detection of homogeneous dwell time EMs (SLOWEM2B, FASTEM2B), any dwell time EM (EM2B), and composite EM where each image fuses more than one dwell time (FUSEDEM2B). We found that FUSEDEM2B, once trained, could be used for all border prediction tasks without compromising quality. The same U-net architecture was also used to train ERRNET, EMINCLUDE, and EMEXCLUDE. We tried the U-net architecture for image homogenization, but achieved better results with conditional GANs.

**Figure S11.**
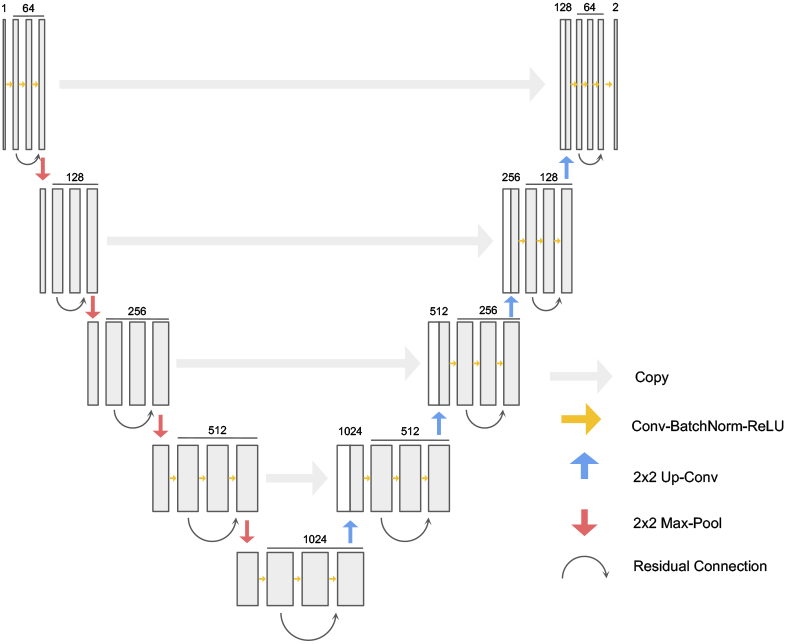
CNN architecture used for the FUSEDEM2B and ERRNET. The architecture is similar to U-Net (Ronneberger et al., 2015), but has 3 layers of (Convolution, Batch–Normalization, ReLU) in each layer and has additional residual connections (He et al. (2016)). The architecture is fully convolutional and for both FUSEDEM2B and ERRNET the input dimension is 1, respectively for the grayscale image and the border probability. In both cases the output dimension is 2, respectively for 0:not-border,1:border and 0:no-error,1:error

### Architecture for FUSEDEM2B and ERRNET

The selected architecture, similar to the U-Net (Ronneberger et al., 2015), shown in **Figure S11** has 3 sets of 2D-Convolution, Batch-Normalization(Ioffe and Szegedy, 2015), ReLU in each layer. We use residual connections (He et al., 2016) adding the output of the first convolution to the last one in each layer. This architecture showed the highest segmentation accuracy when varying the number of CBR (Conv-BatchNorm-ReLU) in each layer (2 ~4), the usage of residual connections, and the type of residual connections (concatenation or addition).

### U-Net architecture for EMEXCLUDE

We trained a fully convolutional U-Net model over 200 epochs, employing a learning rate of 0.01. The model was configured with five layers of depth and filter sizes progressively sequenced as 32, 64, 128, 256, and 512. To introduce non-linearity and manage potential negative inputs, we incorporated a leakyReLU activation function.

### Image Normalization and Augmentation

To train the FUSE-DEM2B network, we used the CLAHE (Pizer et al., 1990) normalization with *clipLimit=3* to bring all images to a common color space. We used on the fly rotation, flip, translation to augment the images in the training set. Although images are naturally 2048 × 1768, we sub-sampled 256 × 256 squares to train the network. To allow the network to deal with images with multiple dwell times, we randomly replace patches at random locations with different dwell times (**Figure S12**). Specifically, each sample was generated by choosing a baseline image at a single dwell time and replacing up to 30 patches with a maximum size of 11 × 11 pixels with the corresponding pixels of an image with longer dwell time.

**Figure S12.**
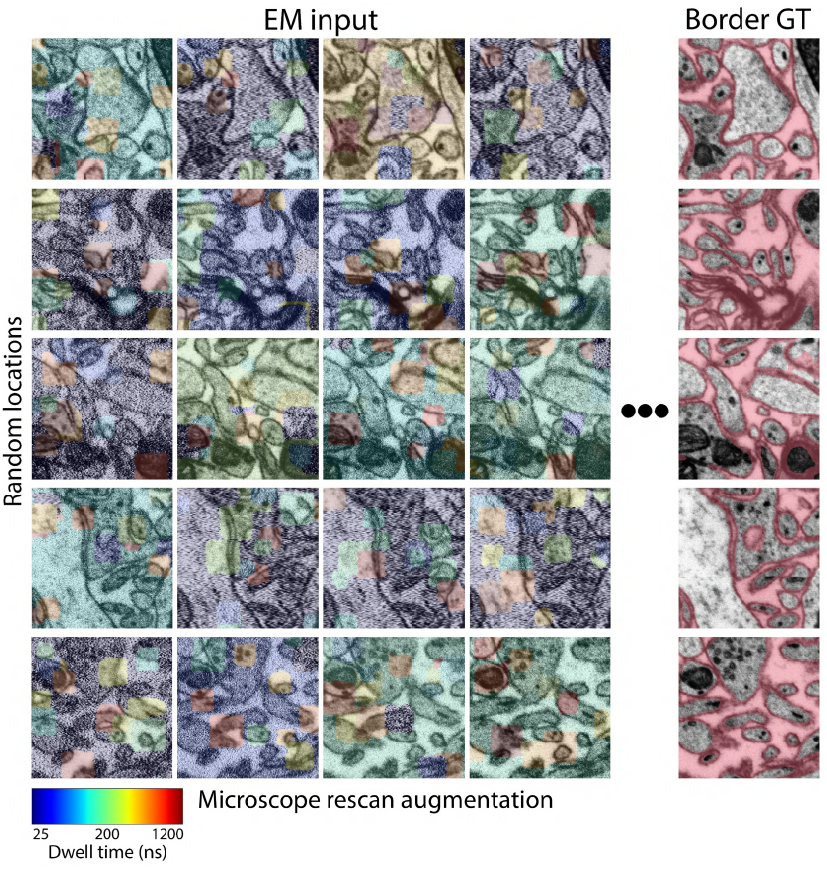
Dwell time rescan data augmentation. Rows 1-5 show different locations in the EM sample. Columns 1-4 show different augmented composite images that were taken at different dwell times; short dwell time pixels in blue, representing 25 ns/pixel scans; long dwell time pixels in red, representing 1200 ns/pixel. Column 5 shows the groundtruth classes for each region that were obtained from the long dwell time neural network (SLOW2EM). The aim of FUSEDEM2B is to classify border pixels. Additional augmentations such as translation, rotation, and flip are used during training.

To train ERRNET, we normalized border probabilities to [0,1] as an input to the network. We used the same procedure for on the fly translation and rotation but did not replace patches.

### Training Procedure

We used the Pytorch framework (Paszke et al., 2019) to implement and optimize the network. The Adam optimizer (Kingma and Ba, 2014) with learning rate 0.001 was used to update the network parameters. We used a batch size of 16 images. We trained the FUSEDEM2B network for 50000 gradient steps. We evaluated validation loss every 1000 steps over 100 batches. The network converged after ~ 35000 gradient steps. The same procedure was used to train ERRNET. ERRNET converged after ~8000 gradient steps.

### Image Translation Networks

IMAGEHOMOGENIZER uses a conditional GAN called pix2pix (Isola et al., 2016), consisting of a generator CNN and discriminator CNN. The generator includes an encoder and decoder that downsamples and then up-samples the input image. The discriminator tries to discriminate between slow EM and translated EM. At the training stage, we use a batch size of 1 and randomly crop 128 × 128 image tiles from a larger composite EM image. The model is first trained with a constant learning rate of 0.0002 for 100 epochs and then for another 100 epochs, during which the learning rate decays to zero. At the inference stage, the whole composite EM image is passed to the model without cropping.

### Image stitching and alignment

The stitching and alignment of the sample volume was performed on composite dwell time images. After applying a band-pass filter to raw images, we used conventional block matching technique to obtain matching points between neighboring images, from which elastic transformations mapping the raw data to the aligned volume were computed by mesh relaxation. Code for stitching and alignment is available at Stitching and alignment code. We applied the same stitching and alignment transformations to the fast, composite, and homogenized images to produce three sets of final volumes.

### Synapse segmentation and neuronal reconstruction Neuron reconstruction technique

To reconstruct neurons in 3D, we applied a lightweight segmentation method that we previously used to reconstruct neurons from the same sample imaged by a multi-beam SEM (Karlupia et al., 2023) and tissue prepared using a whole mouse brain staining technique (Lu et al., 2023). First, pixels straddling intra-cellular spaces were predicted by a CNN, based on the pre-trained FUSEDEM2B network. To improve the network accuracy, we fine-tuned FUSEDEM2B using thirty-six 1024 × 1024 SmartEM tiles obtained from random locations in the target volume and annotated by an expert. Predictions from FUSEDEM2B were used as a starting point for the annotation process of the training set. All sections were segmented in 2D using the fine-tuned network and watersheds (Pavarino et al., 2023). Second, a CNN was trained to predict from the EM the medial axis of all objects in 2D. This process required no additional human annotation. Third, 2D object segments were agglomerated across sections based on shape alignment and similarity. In addition, 2D segments were agglomerated if their medial axes were well-aligned using a fixed threshold determining large overlaps. Fourth, agglomerated objects containing a large number of adjacent 2D segments were flagged as objects with possible merge errors. This was done by building a Regional Adjacency Graph whose nodes are 2D segments and edges represent spatial adjacency. Then these objects were reagglomerated iteratively from the original 2D object segments until the merge-error criterion was attained using an iterative clustering technique (Bailoni et al., 2022). Fifth, orphans were detected and connected to other orphans or non-orphan objects based on their best estimate from the agglomeration graph, i.e., connecting them to objects that did not pass the agglomeration threshold in the first iteration. The results of the reconstruction are shown in **Figure 7C**.

### Criterion for filtering dendritic spines

Three dendrites were randomly selected for quantitative analysis. We defined correctly segmented spines as spines whose segmentation includes their synapse-containing regions. Incorrectly segmented spines were split errors that occurred before the synaptic region. To avoid confusing spines with dendritic filopodia, we excluded putative spines from analysis if no potential synapse was contained in the image volume. There observed three types of error: *Type 1* errors occur when the spine is prematurely truncated by a split error that occurs before the spine’s corresponding synapse that was not due to an obvious image artifact (e.g., tissue preparation, folds in the section). *Type 2* errors occur when the spine is not tracked at all due to a split error at its base on the dendrite that was not due to an obvious image artifact. *Type 3* errors occur when the spine is lost due to an obvious artifact. We observed such errors caused by local aberrations in tissue preparation in sections 56, 65, 66, 77 and 88. The distribution of incorrect spines and their corresponding error type is shown in Table S1. To characterize only errors that might be associated with the SmartEM technique, we exclude the rate of Type 3 errors from consideration.

**Figure S13.**
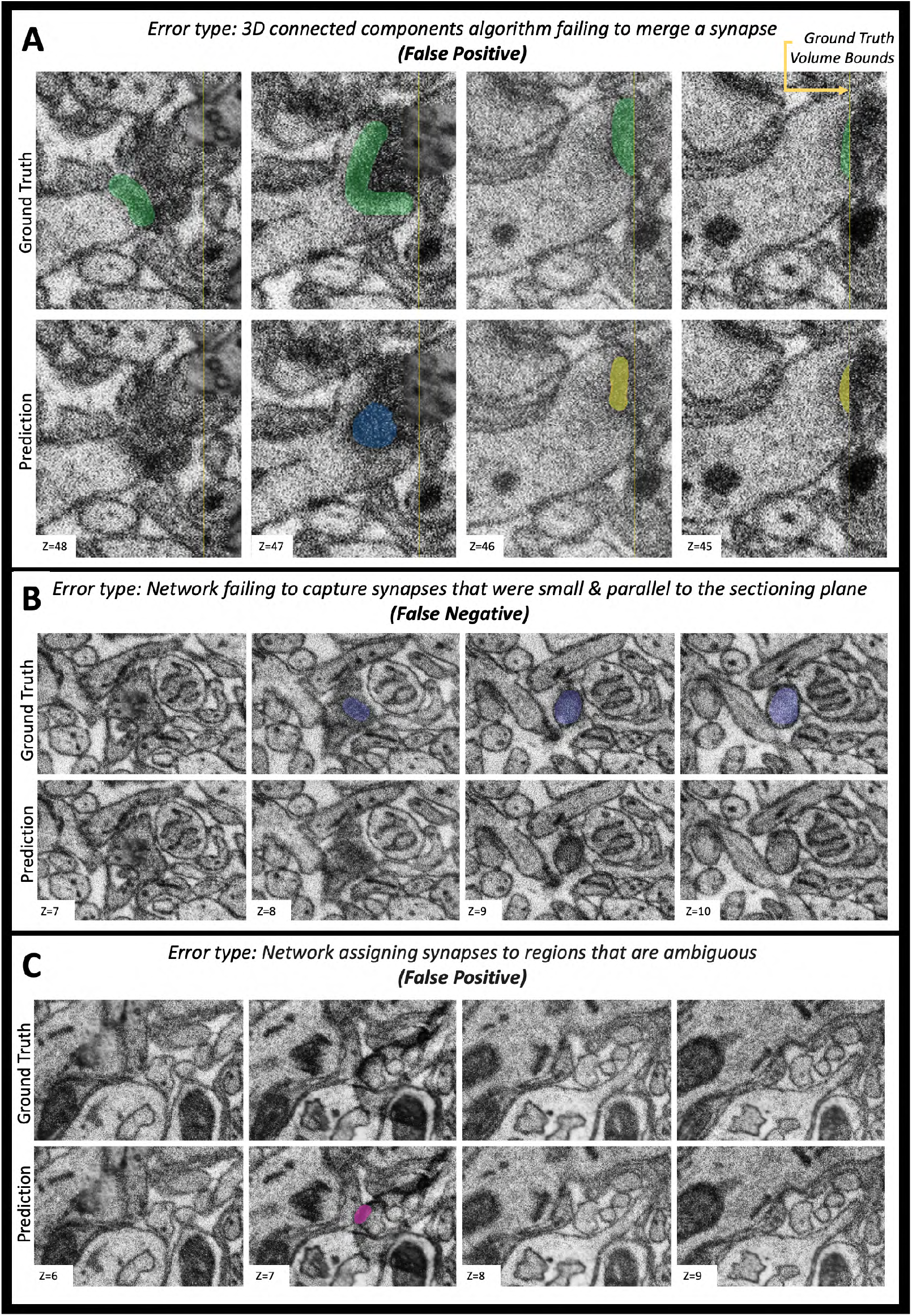
Examples of synapse detection error types. Each example includes synapse ground truth and corresponding model predictions (top and bottom row of each panel respectively), for a span of four images. **A**. False positive error (i.e., synapse identified by the model that is not in the ground truth set) caused by 3D connected components algorithm failing to merge a synapse ([neuroglancer](https://neuroglancer.bossdb.io/#!https%3A%2F%2Fbossdb-neuvue-datalake.s3.amazonaws.com/public/states/dc6319d0-96db-407a-9e42-05f995567c67.json)). **B**. False negative error (i.e., synapse not identified by the model that is in the ground truth set) caused by the network failing to capture synapses that were small and parallel to the sectioning plane ([neuroglancer](https://neuroglancer.bossdb.io/#!https%3A%2F%2Fbossdb-neuvue-datalake.s3.amazonaws.com/public/states/dc6319d0-96db-407a-9e42-05f995567c67.json)). **C**. False positive error resulting from the network assigning synapses to regions that are ambiguous ([neuroglancer](https://neuroglancer.bossdb.io/#!https%3A%2F%2Fbossdb-neuvue-datalake.s3.amazonaws.com/public/states/dc6319d0-96db-407a-9e42-05f995567c67.json)).

### Synapse segmentation technique

Synaptic active zones were manually segmented in VAST (Berger et al., 2018) and agreed upon by two expert annotators. Two ground truth volumes, GT1 and GT2, were generated of sizes 7 × 3.5 × 3 *µ*m^3^ and 4 × 4 × 3 *µ*m^3^ respectively. A U-Net was trained on GT1 to predict active zones from EM images via the PyTorch Connectomics library (Lin et al., 2021). To avoid edge effects, the trained network was applied on a padded version of the EM from GT2. A threshold of 0.8 was applied to the outputs of this network, followed by 3D connected components with 26-connectivity using the cc3d library (Silversmith, 2021). We removed segments that were smaller than 400 voxels. All parameters for post-processing were determined without studying the statistics of GT2; the voxel threshold was obtained by rounding down the smallest segment size in GT1. The results were finally cropped to account for the fact that the EM input was padded.

### Synapse validation

To assess synapse segmentability, we replicated CONFIRMS (Bishop et al., 2021), a quantification tool developed for EM pipeline validation. In short, synapse segments were converted into keypoints by determining the location of their centroid and matched according to the distances between these keypoints. The matches were manually verified in [neuroglancer](https://neuroglancer.bossdb.io/#!https%3A%2F%2Fbossdb-neuvue-datalake.s3.amazonaws.com/public/states/dc6319d0-96db-407a-9e42-05f995567c67.json). When the matching algorithm incorrectly assigns a synapse a certain label (ex: synapse is assigned as false positive, when it is really a true positive), it was corrected by experts. We conservatively made corrections to the results of the matching algorithm. For example, if expert annotators see a false positive but believe it to be an ambiguous synapse, it was still treated as a false positive. The precision, recall and F1-Scores were calculated after these corrections were made.

### Examples of errors in synapse reconstruction

All errors were manually inspected by experts in [neuroglancer](https://neuroglancer.bossdb.io/#!https%3A%2F%2Fbossdb-neuvue-datalake.s3.amazonaws.com/public/states/dc6319d0-96db-407a-9e42-05f995567c67.json). These errors could be classified into three main categories. The first error type was caused by the 3D connected components algorithm failing to merge a synapse, resulting in two synapses (**Figure S13A**, [neuroglancer](https://neuroglancer.bossdb.io/#!https%3A%2F%2Fbossdb-neuvue-datalake.s3.amazonaws.com/public/states/dc6319d0-96db-407a-9e42-05f995567c67.json)). The second error type was a result of the network failing to capture synapses that were small and parallel to the sectioning plane (**Figure S13B**, [neuroglancer](https://neuroglancer.bossdb.io/#!https%3A%2F%2Fbossdb-neuvue-datalake.s3.amazonaws.com/public/states/dc6319d0-96db-407a-9e42-05f995567c67.json)). The third main error type was the network assigning synapses to regions that are ambiguous (**Figure S13C**, [neuroglancer](https://neuroglancer.bossdb.io/#!https%3A%2F%2Fbossdb-neuvue-datalake.s3.amazonaws.com/public/states/dc6319d0-96db-407a-9e42-05f995567c67.json)). We observed no clear correlation between regions of rescanned EM and errors in synapse reconstruction.

### Distribution of synapse sizes

We verified that synapses in SmartEM volumes can be reconstructed with accurate morphology in 3D. To do so, we first visualized the cumulative distributions of synapse size for the network predictions, the true positive predictions, and the ground truth (**Figure S14A**). To further investigate how well the synapse sizes of true positive predictions match ground truth, we plotted the correlation in synapse size between matching objects in these volumes (**Figure S14B**). We calculated the Pearson correlation on the logarithmically scaled synapses (*r* = 0.91, *p* = 1.51 × 10^−38^), implying that the automated reconstruction of synapses in SmartEM is a strong predictor of the real morphology in 3D. Nevertheless, proofreading of synapses is required, as is seen by the small number of smaller objects that diverge from the trend. We also plotted the performance of the network (precision, recall and the F1-score) conditioned on the synapse size (**Figure S14C**). We observed that the performance of the network improves with the size of the synapse, consistent with our intuition that larger synapses are easier to detect.

### Statistical tests

All statistical tests were done using the Wilcoxon signed-rank test for paired samples. The test was used to assess the cases where two dwell times produce similar segmentation quality by comparing the variation of information of individual samples to a single reference taken at a longer dwell time.

### Sample Preparation

The preparation of the mouse cortex is described in (Karlupia et al., 2023) and the human cortex in (Shapson-Coe et al., 2024). The worm was prepared as follows. Several *C. elegans* males were transferred from a mixed-culture plate (N2 wildtype strain) to a separate plate seeded with *E. coli* OP50, where they were kept for 16h prior to high pressure freezing (HPF). L4 larvae were selected, and they all became adults by the time of HPF. For HPF, we used gold-coated copper carriers (16770152 and 16770153, Leica), which were soaked in a 2% lecithin in chloroform solution and allowed to dry to render their surface non-stick (Mulcahy et al., 2018). Live *C. elegans* males were transferred from the culture plate to the carrier together with a small amount of *E. coli* substrate. The samples were then frozen using a high pressure freezer (EM ICE, Leica). This was followed by freeze-substitution, which was carried out in a programmable unit (EM AFS2, Leica), using a 1% ddH2O, 1% OsO4, and 1% glutaraldehyde in acetone solution, at −90C for 48h, after which the temperature was increased by 5C per hour until it reached 20C (Weir et al., 2020). The sample pellets were then washed with acetone (3 times) and infiltrated with 50% Epon in acetone for 1 h, 75% Epon in acetone overnight, 100% Epon for 1 hour (the last step was repeated twice), and finally cured at 60C. The samples were imaged with microCT to check for major cracks. 35 nm sections were cut and collected on kapton tape using a Leica EM UC6 ultramicrotome and ATUM section collecting device (Kasthuri et al., 2015; Baena et al., 2019). The tape with the sections was mounted on silicon wafers and the sections were then post-stained with uranyl acetate and lead citrate as described in Baena et al. (2019). The samples were kept under vacuum for at least 24h before imaging to minimize any beam-related damage due to residual water.

**Figure S14.**
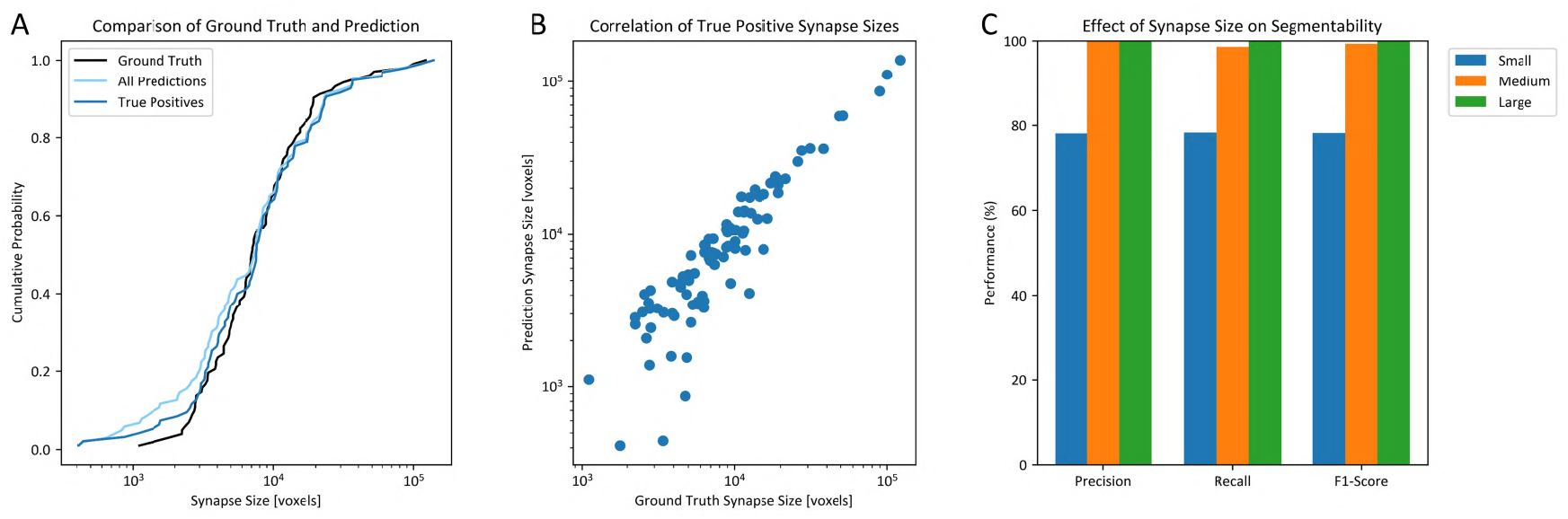
**A**. The cumulative distribution of synapse sizes for the network predictions, the true positive network predictions and ground truth. **B**. Correlation plot of synapse sizes between the true positive network prediction and ground truth (Pearson correlation: *r* = 0.91, *p* = 1.51 × 10^−38^). **C**. Precision, recall and F1-score of network conditioned on synapse size (small: 400 – 4000 voxels, medium: 4000 – 40000 voxels, large: > 40000 voxels).

### Visualization and proofreading tools

All SmartEM datasets and evaluation outputs are publicly available through BossDB (The Brain Observatory Storage Service and Database) (Hider et al., 2022), a cloud-native platform built for scalable storage, visualization, and dissemination of large-scale neuroimaging and connectomics data. The SmartEM data collection includes image volumes acquired using multiple acquisition modalities – including SmartEM and the corresponding short scan – for cross-modal benchmarking and assessment. A manifest of available datasets and derived products is provided on the BossDB project page (https://bossdb.org/projects). Raw image stacks, alignment results, and segmentation outputs were processed using containerized workflows deployed in a cloud-based infrastructure to ensure reproducibility and scalability. SmartEM volumetric data are stored in Google’s precomputed format (Maitin-Shepard et al., 2021) for ease of integration and real-time visualization within the BossDB ecosystem.

Visualization and human-in-the-loop evaluation were conducted using open-source tools, including Neuroglancer (Maitin-Shepard et al., 2021) and NeuVue (Xenes et al., 2022). Neuroglancer provided an interactive, web-based interface for exploring high-resolution volumetric data, while NeuVue supported structured task queues for annotation and proofreading of EM volumes.

**Figure S15.**
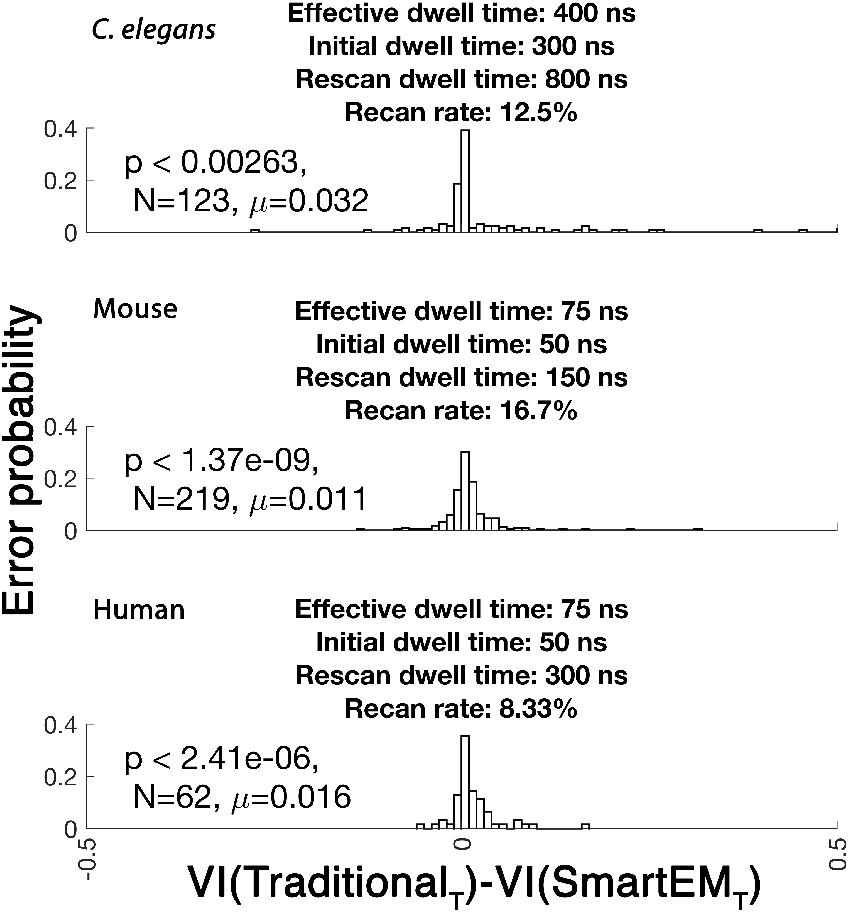
Distribution of segmentation error differences (standard minus SmartEM) in three datasets-the *C. elegans* nerve ring (top), mouse cortex (middle), and human temporal lobe (bottom)—collected at the same time-matched effective dwell time. Parameter settings for each case (effective dwell times, initial dwell times, rescan dwell times, and rescan rates) are shown in the panel titles. Each distribution plots the Variation of Information (VI) error from the standard pipeline minus that from SmartEM for a collection of N images; Positive values indicate lower error in SmartEM. All three datasets exhibit a right-tailed distribution and a significantly higher mean error in the standard pipeline (mean differences are shown). Sign tests further confirm that the standard pipeline consistently produces larger segmentation errors than SmartEM (p-values are indicated for each dataset).

**Table S1.**
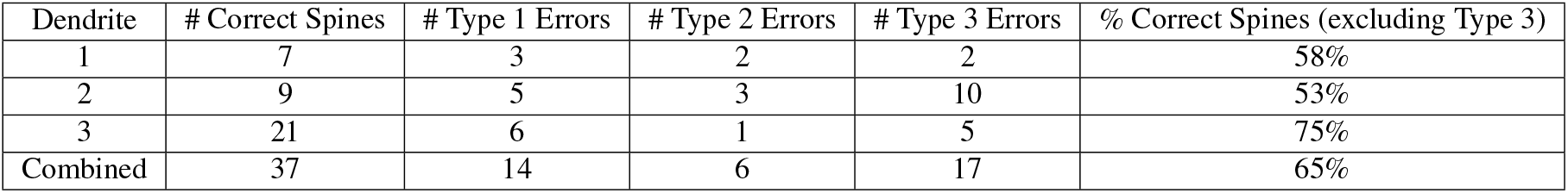
Distribution of correctly and incorrectly segmented dendritic spines by automated reconstruction.

| Dendrite | # Correct Spines | # Type 1 Errors | # Type 2 Errors | # Type 3 Errors | % Correct Spines (excluding Type 3) |
| --- | --- | --- | --- | --- | --- |
| 1 | 7 | 3 | 2 | 2 | 58% |
| 2 | 9 | 5 | 3 | 10 | 53% |
| 3 | 21 | 6 | 1 | 5 | 75% |
| Combined | 37 | 14 | 6 | 17 | 65% |

